# Species-specific sensitivity to TGFβ signaling and changes to the Mmp13 promoter underlie avian jaw development and evolution

**DOI:** 10.1101/2020.12.23.424223

**Authors:** Spenser S. Smith, Daniel Chu, Tiange Qu, Richard A. Schneider

## Abstract

Developmental control of jaw length is critical for survival. The jaw skeleton arises from neural crest mesenchyme and previously we demonstrated that these progenitors upregulate bone-resorbing enzymes including *Matrix metalloproteinase 13* (*Mmp13)* when generating short quail beaks versus long duck bills. Inhibiting bone resorption or *Mmp13* increases jaw length. Here, we uncover mechanisms establishing species-specific levels of *Mmp13* and bone resorption. Quail show greater activation of, and sensitivity to *Transforming Growth Factor-Beta* (*TGFβ*) signaling than duck; where mediators like SMADs and targets like *Runx2,* which bind *Mmp13*, become elevated. Inhibiting TGFβ signaling decreases bone resorption. We discover a SMAD binding element in the quail *Mmp13* promoter not found in duck and single nucleotide polymorphisms (SNPs) near a RUNX2 binding element that affect expression. Switching the SNPs and SMAD site abolishes TGFβ-sensitivity in the quail *Mmp13* promoter but makes duck responsive. Thus, differential regulation of TGFβ signaling and *Mmp13* promoter structure underlie avian jaw development and evolution.

## INTRODUCTION

Jaws are among the most highly adapted and modified structures of vertebrates, and they facilitate complex behaviors like feeding, respiration, and vocalization. In this context, precise developmental regulation of jaw length is crucial for survival (Schneider, 2015). By comparing jaw development between white Pekin duck and Japanese quail, we have shown in prior work that neural crest mesenchyme (NCM), which is the progenitor population that gives rise to the jaw skeleton, employs a variety of stage- and species-specific mechanisms to govern jaw length through successive phases of development (Jheon and Schneider, 2009; Fish and Schneider, 2014b; Schneider, 2018b; Schneider, 2018a). Quail have much shorter jaws compared to those of duck, and during the early migration of NCM from the anterior neural tube, duck embryos allocate more progenitors to the presumptive jaw region (Fish et al., 2014). Once these NCM populations arrive, their growth trajectories further diverge due to autonomous molecular programs for proliferation and differentiation that are tied to intrinsic rates of maturation and species-specific regulation of multiple signaling pathways (Eames and Schneider, 2008; Merrill et al., 2008; Mitgutsch et al., 2011; Hall et al., 2014; Ealba et al., 2015).

A key finding from these studies is the identification of a previously unrecognized developmental mechanism governing jaw length, which is NCM-mediated bone resorption. Quail have higher levels of bone resorption enzymes than do duck during late stages of jaw development (Ealba et al., 2015), including tartrate-resistant acid phosphatase (TRAP) and *Matrix metalloproteinase 13 (Mmp13*), which is a bone-resorbing collagenase secreted by osteocytes and other cells (Johansson et al., 1997; Sasano et al., 2002; Behonick et al., 2007; Chen et al., 2012a). Osteocytes regulate local bone resorption through a process called perilacunar remodeling (Belanger, 1969; Hatori et al., 2004; Inoue et al., 2006; O’Brien et al., 2008; Qing et al., 2012; Tang et al., 2012; Xiong et al., 2014; Dole et al., 2017; Alemi et al., 2018) and in the jaw skeleton osteocytes are derived exclusively from NCM (Le Lièvre, 1978; Noden, 1978; Helms and Schneider, 2003). Transplanting presumptive cephalic NCM from quail to duck dramatically elevates expression of bone resorption enzymes and generates chimeras with shorter quail-like jaws, whereas blocking bone resorption using a bisphosphonate or an MMP13 inhibitor significantly lengthens the jaw (Ealba et al., 2015). Additionally, knockdown of *Mmp13* alters jaw growth in zebrafish (Hillegass et al., 2007) and affects the shape of craniofacial structures during tadpole development (Pinet et al., 2019). Patients with mutations in *Mmp13* also display jaw size defects (Kennedy et al., 2005). Taken together, such experiments reveal that NCM controls bone resorption and that there is a link between bone resorption, *Mmp13* activity, and jaw length. However, what has remained unclear are the molecular mechanisms that lead to differential regulation of *Mmp13* and the species-specific control of bone resorption in relation to jaw length.

To address this question, in the current study we focus on the *Transforming Growth Factor Beta* (TGFβ) signaling pathway, which is known to mediate bone deposition and resorption, as well as *Mmp13* expression (Stouffer and Owens, 1994; Moses and Serra, 1996; Vinals and Pouyssegur, 2001; Kim et al., 2004; Selvamurugan et al., 2004a; Fang et al., 2012; Crane and Cao, 2014; Wu et al., 2016). TGFβ ligands interact at the plasma membrane with a receptor dimer consisting of type I and type II TGFβ receptors. Upon binding of TGFβ ligand, the type II receptor (TGFβR2) transphosphorylates and activates the type I receptor (TGFβR1), initiating an intracellular signaling cascade involving phosphorylation of SMAD2 and SMAD3. These activated SMADs form a complex with SMAD4, allowing translocation into the nucleus and interaction with SMAD binding elements and other DNA binding proteins to activate or repress transcription of target genes (Heldin et al., 1997; Dennler et al., 1998; Derynck et al., 1998; Li et al., 1998; Massague and Wotton, 2000; Alliston et al., 2001; Derynck and Zhang, 2003). Inhibiting TGFβ receptor kinases suppresses *Mmp13* expression *in vivo* (Dunn et al., 2009), while normal TGFβR1 activity positively affects MMP13 and the perilacunar remodeling of bone (Dole et al., 2017). Similarly, other target genes such as *Runx2*, which is a major transcription factor expressed by osteoblasts (Ducy et al., 1997; Komori et al., 1997; Karsenty et al., 1999; Selvamurugan et al., 2004a; Derynck et al., 2008), can be induced or repressed by TGFβ ligands depending on the levels of exposure and the complement of transcriptional co-factors (Lee et al., 2000; Alliston et al., 2001; Selvamurugan et al., 2004b; Wu et al., 2016). Overexpressing *Runx2* in NCM can shorten the jaw (Hall et al., 2014), whereas patients with a *Runx2* haploinsufficiency can develop an enlarged lower jaw (Gorlin et al., 1990; Jaruga et al., 2016; Pan et al., 2017).

To identify mechanisms that control the differential regulation of *Mmp13* and potentially link bone resorption and jaw length, we assay for species-specific expression of ligands, receptors, and effectors of the TGFβ pathway in chick, quail, and duck embryos during key stages of jaw development when bone is being deposited and resorbed. We employ these three birds for several reasons. First, for comparative studies, a three-taxon analysis in which two taxa are more closely related than either are to a third taxon, is generally accepted as a robust strategy for making the most parsimonious inferences about evolution (Nelson and Platnick, 1991; Mavrodiev et al., 2019; Rineau et al., 2020). As Galliformes, chick and quail are more closely related to each other phylogenetically (separated by around 50 million years) and they diverged from a common ancestor with Anseriformes, which include duck, over 100 million years ago (Pereira and Baker, 2006; Hackett et al., 2008; Kan et al., 2010). Second, chick and quail are more similar in terms of their jaw shape when compared to duck, whereas the chick jaw is more similar to duck in terms of size (Schneider and Helms, 2003; Eames and Schneider, 2008; Mitgutsch et al., 2011; Smith et al., 2015). Third, chick is a long-established experimental model system with a well annotated genome (Stern, 2005; Lwigale and Schneider, 2008; Sauka-Spengler and Barembaum, 2008; Jheon and Schneider, 2009; Fish and Schneider, 2014a; Abramyan and Richman, 2018; Gammill et al., 2019), which helps in analyzing data from quail and duck (Ealba and Schneider, 2013; Chu et al., 2020).

We quantify expression of TGFβ pathway members and observe some of the highest levels in quail versus duck and chick. MMP13 protein is also elevated in domains that overlap directly with TRAP staining in jaw bones. To test if the species-specific levels of TGFβ pathway activation are due to intrinsic differences in sensitivity to TGFβ signaling, we perform cell and organ culture experiments. We treat chick and duck cells, as well as lower jaw primordia from chick, quail, and duck with recombinant (r) TGFβ1. These experiments reveal species-specific differences in sensitivity to rTGFβ1, with downstream targets more upregulated in quail than in duck. We also find that inhibiting TGFβR1, SMAD3, or MMP13 reduces bone resorption in the developing jaw.

To identify molecular mechanisms underlying species-specific differences in sensitivity to TGFβ signaling and in the response of downstream targets, we examine the structure and function of the *Mmp13* promoter in chick, quail, and duck. *Mmp13* is likewise a target of RUNX2 and its promoter contains RUNX2, SMAD, and other binding elements that regulate its expression (Pendas et al., 1997; Selvamurugan et al., 2004a; Selvamurugan et al., 2004b; Wang et al., 2004; Selvamurugan et al., 2006; Selvamurugan et al., 2009; Chen et al., 2012a; Meyer et al., 2016; Takahashi et al., 2017; Arumugam et al., 2018; Young et al., 2019). Moreover, genetic polymorphisms in the *Mmp13* promoter have been shown to affect transcriptional activity and phenotypic variation in chick (Yuan et al., 2016) and humans (Ye, 2000; Benderdour et al., 2002; Yoon et al., 2002; Yan and Boyd, 2007; Achari et al., 2008; Hashimoto et al., 2013). We discover that species-specific expression is mediated by the structure of the *Mmp13* promoter, ostensibly at RUNX2 and SMAD binding elements. We generate species-specific reporter constructs with or without these binding elements and find that quail and chick promoters show high activity whereas the duck promoter shows low to no activity. Removing these binding elements reduces activity with quail and chick promoters but adding them increases activity with the duck promoter. We also identify two single nucleotide polymorphisms (SNPs) directly downstream of the RUNX2 binding element that distinguish quail and chick from duck. Switching these SNPs between species reduces activity in quail and chick promoters but increases activity in duck. Overall, our results indicate that multiple levels of gene regulation in the TGFβ signaling pathway mediate *Mmp13* expression, bone resorption, and ultimately species-specific variation in jaw length.

## MATERIALS AND METHODS

### The Use of Avian Embryos

Fertilized eggs of chicken (*Gallus gallus*), Japanese quail (*Coturnix coturnix japonica*), and white Pekin duck (*Anas platyrhynchos domestica*) were purchased from AA Lab Eggs (Westminster, CA) and incubated at 37.5°C in a humidified chamber (GQF Hova-Bator 1588, Savannah, GA) until they reached embryonic stages appropriate for analyses. For all experiments, we adhered to accepted practices for the humane treatment of avian embryos as described in S3.4.4 of the AVMA Guidelines for the Euthanasia of Animals: 2013 Edition (Leary et al., 2013). Embryos were matched at equivalent stages the using the Hamburger and Hamilton (HH) staging system, a well-established standard that utilizes an approach based on external morphological characters, that is independent of body size and incubation time, and that can be adapted to other avian species such as quail and duck (Hamburger and Hamilton, 1951; Hamilton, 1965; Ricklefs and Starck, 1998; Starck and Ricklefs, 1998; Schneider and Helms, 2003; Lwigale and Schneider, 2008; Jheon and Schneider, 2009; Ainsworth et al., 2010; Mitgutsch et al., 2011; Fish and Schneider, 2014a; Smith et al., 2015).

### Histological Staining and Immunohistochemistry

Chick, quail, and duck embryos were collected at HH40 in 4% paraformaldehyde (PFA) (15714, Electron Microscopy Sciences, Hatfield, PA, USA) overnight at 4°C (Schneider, 1999; Schneider et al., 2001). To detect tartrate-resistant acid phosphatase (TRAP) in whole mount, chick, quail, and duck jaws were stained using the Acid Phosphatase Leukocyte kit (387A-1KT, MilliporeSigma, Burlington, MA, USA) following the manufacturer’s protocol, except 7 mg/mL Fast Red Violet (F3381, MilliporeSigma, Burlington, MA, USA) were used in place of the Fast Garnet GBC Base Solution (Ealba et al., 2015). To detect bone deposition and resorption in sections, embryos were dehydrated in methanol, embedded in paraffin, and cut into 10 µm sagittal sections. Sections were deparaffinized, rehydrated, and adjacent sections were stained with Milligan’s trichrome at room temperature as previously described (Presnell and Schreibman, 1997; Schneider et al., 2001; Eames and Schneider, 2005; Tokita and Schneider, 2009; Solem et al., 2011; Hall et al., 2014) or for TRAP at 37°C for 1 hour (Ealba et al., 2015).

Immunohistochemistry (IHC) was performed on adjacent sections. For antigen retrieval, sections were heated in a microwave to 95°C in 10 mM sodium citrate buffer for 10 min and endogenous peroxidase activity was blocked with 3% hydrogen peroxide for 15 minutes. Sections were incubated with 1 µg/ml of a custom-made MMP13 rabbit polyclonal primary antibody (GenScript, Piscataway, NJ, USA; Supplemental Table S1) overnight at 4°C. Sections were labeled with 1:500 goat anti-rabbit Alexa Fluor 647 secondary antibody (A32733, Thermo Fisher Scientific, Waltham, MA, USA) overnight at 4°C. 10 mg/ml Hoechst 33342 dye (62249, Thermo Fisher Scientific, Waltham, MA, USA) was used to stain nuclei. Lower jaw sections were imaged using a Nikon AZ100 C2 macroconfocal microscope and image acquisition system (Nikon Instrument, Inc., Melville, NY) for IHC, and a Leica DM 2500 (Leica Microsystems, Inc. Buffalo Grove, IL) with a color digital camera system (SPOT Insight 4 Megapixel CCD, Diagnostic Instruments, Inc., Sterling Heights, MI) for Trichrome and TRAP.

To highlight specific-specific differences in jaw morphology, chick, quail, and duck heads were fixed in 4% PFA overnight and stained for 20 minutes with 0.02% ethidium bromide (1610433, Bio-Rad, Hercules, CA, USA) using a previously published protocol (Eames and Schneider, 2005). Samples were washed 3 times in 1X PBS. To detect TRAP in whole mount, lower jaws were skinned, fixed in 4% PFA, and stained for 1.5 hours at 37°C. Samples were then cleared in glycerol (525342C, Thermo Fisher Scientific, Waltham, MA, USA). Ethidium bromide-stained and TRAP-stained samples were imaged on a dissecting microscope (Leica MZFLIII) using either epifluorescent, transmitted, and/or incident illumination and a color digital camera system (SPOT Insight 4 MP).

### Quantitative PCR

Lower jaws were dissected from chick, quail, and duck embryos and total RNA was extracted using the RNeasy Plus Mini Kit (74136, Qiagen, Hilden, Germany) following the manufacturer’s protocol. Lower jaws were resuspended in 600 μl of RTL plus buffer supplemented with 1% β-mercaptoethanol (M3148-100ML, MilliporeSigma, Burlington, MA, USA) and Reagent DX (19088, Qiagen, Hilden, Germany). HH31 and HH34 lower jaws were processed in a Bead Mill 24 Homogenizer (15-340-163, Fisher Scientific, Waltham, MA, USA) at 5 m/s for 30 s with 1.4 mm ceramic beads (15-340-153, Fisher Scientific, Waltham, MA, USA). HH37 and older lower jaws were homogenized at 5 m/s for 60 s with 2.8 mm ceramic beads (15-340-154, Fisher Scientific, Waltham, MA, USA). Following purification of total RNA, residual genomic DNA was removed using TURBO DNA-free Kit (AM1907, Invitrogen, Carlsbad, CA, USA). DNased RNA was reverse-transcribed using iSCRIPT (1708841, Bio-Rad). Gene expression was analyzed by quantitative PCR (qPCR) with iQ SYBR Green Supermix (1708882, Bio-Rad, Hercules, CA, USA) and normalized to 18S rRNA following previously published protocols (Dole et al., 2015; Smith et al., 2016). Primers were designed (Geneious Prime, Version 2020.2.4) to amplify conserved regions among chick, quail, and duck for members and targets of the TGFβ pathway including *Tgfβ1*, *Tgfβ2*, *Tgfβ3*, *Tgfβr1*, *Tgfβr2, Tgfβr3, Activin a receptor like type 1 (Acvrl1), Smad2, Smad3, Runx2, Mmp2, Mmp9, Mmp13, Mmp14, Plasminogen activator inhibitor 1 (Pai1), Collagen type 1 α 1 (Col1a1), Osteocalcin (Ocn), Cathepsin K (Ctsk),* and *Sclerostin (Sost)* (Supplemental Table S2). Criteria for experimental design included limiting primers to 20 bp in length, amplifying regions of ∼150 bp, using an annealing temperature of 60°C, keeping GC content around 50%, minimizing self-complementarity (*i.e.,* primer-dimers), and amplifying regions that span exon-exon junctions. To account for alternative efficiencies of primer binding between species, data were normalized using serial dilutions of pooled cDNA and a standard curve method (Ealba and Schneider, 2013; Dole et al., 2015; Smith et al., 2016). Each sample was assayed in technical duplicate.

### Western Blots

Lower jaws were lysed with 1X RIPA lysis buffer (20-188, MilliporeSigma, Burlington, MA, USA) containing Halt protease inhibitors (78430, Thermo Fisher Scientific, Waltham, MA, USA). A BCA assay (23225, Thermo Fisher Scientific, Waltham, MA, USA) was performed to quantify protein using a SpectraMax M5 microplate reader (Molecular Devices, San Jose, CA, USA). 40 µg of protein was electrophoresed on a 10% SDS polyacrylamide gel as previously described (Smith et al., 2016). Proteins were transferred to an Immobilon-P PVDF membrane (IPVH00010, MilliporeSigma, Burlington, MA, USA). Membranes were probed with 1:1000 rabbit anti-human pSer423/pSer425 SMAD3 antibody (NBP1-77836, Novus Biologicals, Littleton, CO, USA), 1:1000 rabbit anti-human SMAD3 antibody (NB100-56479, Novus Biologicals, Littleton, CO, USA), 1 µg/ml rabbit anti-chick MMP13 custom-made primary antibody (GenScript, Piscataway, NJ, USA; Supplemental Table 2), 1:1000 rabbit anti-human MMP2 antibody (NB200-193, Novus Biologicals, Littleton, CO, USA), 1:4000 mouse anti-human β-actin antibody (NB600-501, Novus Biologicals, Littleton, CO, USA), 1:15000 goat anti-rabbit IRDye 800CW (925-32211, LI-COR, Lincoln, NE, USA), and 1:15000 donkey anti-mouse IRDye 680RD antibody (925-68072, LI-COR, Lincoln, NE, USA). Fluorescent signal was detected using the Odyssey Imaging System (LI-COR, Lincoln, NE, USA,). Quantifications of protein bands were performed using Image Studio Lite. Protein levels were normalized to β-actin.

### Culture Experiments and TGFβ Pathway Manipulation

For *in vitro* experiments, an embryonic chick fibroblast cell line (DF-1, CRL-12203, ATCC, Manassas, VA, USA) and an embryonic duck fibroblast cell line (CCL-141, ATCC, Manassas, VA, USA) were cultured in complete media, Dulbecco’s Modified Eagle’s Medium (DMEM, 10-013-CV, Corning, Corning, NY, USA) for chick fibroblasts (*i.e.*, DF-1 cells) or MEMα (Thermo Fisher Scientific, Waltham, MA, USA, A10490-01) for duck fibroblasts (*i.e.*, CCL-141 cells) supplemented with 10% Fetal bovine serum (FBS, 97068-085, Lot# 283K18, VWR, Radnor, PA, USA) and 1X penicillin-streptomycin (15140122, Thermo Fisher Scientific, Waltham, MA, USA). Cells were screened monthly for mycoplasma contamination. Cells were plated; serum-deprived for 12 hours; treated with 5 ng/ml recombinant (r) human TGFβ1 derived from HEK293 cells (100-21, PeproTech, Rocky Hill, NJ, USA) for 1, 3, 6, or 24 hours; and then harvested for mRNA and protein analysis. For luciferase assays, cells were treated with 5 ng/ml rTGFβ1 for 24 hours. To disrupt the TGFβ pathway, cells were treated with 1 µM SB431542 (S4317, MilliporeSigma, Burlington, MA, USA) to inhibit TGFβR1 activity or 3µM SIS3 (5291, Tocris/R&D Systems, Minneapolis, MN, USA) to inhibit SMAD3 solubilized in 50 mM DMSO, alone or in combination with rTGFβ1 for 24 hours.

For *ex vivo* experiments, HH34 lower jaws from chick, quail, and duck were dissected, placed on a 0.45 µm membrane filter (HAWP01300, MilliporeSigma, Burlington, MA, USA), cultured for 24 hours in 6-well transwell inserts (10769-192, VWR, Radnor, PA, USA) in complete media, switched to media supplemented with 1% FBS and 1X penicillin-streptomycin, and treated with 10, 25, or 50 ng/ml rTGFβ1 in the media for 6 or 24 hours. Lower jaws were then collected for mRNA and protein analyses. To disrupt the TGFβ pathway, lower jaws from quail and duck were harvested at HH35. Affigel Blue Beads (1537301, 250-300 µm diameter, 50-100 mesh, Bio-Rad, Hercules, CA, USA) were washed with PBS and soaked in either 10 mM SB431542, 10 mM SIS3, 1 mg/ml MMP13 inhibitor (444283, MilliporeSigma, Burlington, MA, USA), 160 µg/ml rTGFβ1, or 100 µg/ml rMMP13 (4442875, MilliporeSigma, Burlington, MA, USA) for 1 hour at room temperature. Concentrations were based on those used previously (Ealba et al., 2015; Havis et al., 2016; Woronowicz et al., 2018). Controls beads were soaked in 10% DMSO for SB431542, SIS3, and MMP13 inhibitor treatments. Control and treatment beads were surgically inserted into the left and right sides (respectively) using forceps, and lower jaws were placed on a 0.45 µm membrane filter, put in transwell inserts, cultured in complete media supplemented with 50 µg/ml ascorbic acid (A61-25, Thermo Fisher Scientific, Waltham, MA, USA) and 10 mM β-glycerol phosphate (AC410991000, Thermo Fisher Scientific, Waltham, MA, USA) for 5 days, and collected for whole mount TRAP staining.

### Quantification of TRAP Staining

To quantify TRAP staining, images of lower jaws treated with TGFβR1, SMAD3, or MMP13 inhibitors were adjusted in Adobe Photoshop 2020 (version 21.2.2) to normalize for exposure, brightness, contrast, saturation, and color balance across samples. Exclusion criteria comprised samples where control and/or treatment beads had fallen out or were misplaced, samples that became substantially malformed or stunted during culture, and samples that were uniformly over- or under-stained with TRAP. The Rectangular Marquee Tool was used in Photoshop to define a 1 mm square area (200 by 200 pixels) centered around either the control or treatment bead on each side of the lower jaw. Images were cropped to the 1 mm square. The Elliptical Marquee Tool was used to delete an equally sized area that covered the beads in each pair of cropped images (i.e., control versus treated sides of the same sample). Cropped images were opened in ImageJ (Fiji Version 2.1.0/1.53g) (Schindelin et al., 2012; Schneider et al., 2012). Cropped images were adjusted using the Color Threshold tool and the Default method so that the same thresholding value was applied to the control and treated sides of each pair. Thresholded images were analyzed using the Analyze Particles function and results were saved as % Area to represent the total amount of TRAP-positive staining (Sawyer et al., 2003; Holland et al., 2019; Mira-Pascual et al., 2020).

### Mmp13 Promoter Sequencing

The *Mmp13* promoters for chick, quail, and duck were sequenced through inverse PCR, which enables amplification of unknown sequences that flank a known region (Ochman et al., 1988; Green and Sambrook, 2019). Genomic DNA for inverse PCR was extracted from embryonic chick, quail, and duck tissues using the Purelink Genomic DNA mini kit (K1820-01, Invitrogen, Carlsbad, CA, USA) following the manufacturer’s protocol. The sequences for exon 1 of chick, quail, and duck were used as the anchor for designing primers and determining restriction sites. Genomic DNA for the inverse PCR was digested with EcoRI-HF (R3101S, NEB, Ipswich, MA, USA). Digested genomic DNA was purified with GeneJET PCR Purification Kit (K0702, Thermo Fisher Scientific, Waltham, MA, USA) and then ligated with Rapid DNA Ligation Kit (K1422, Thermo Fisher Scientific, Waltham, MA, USA). Inverse PCR was performed on the ligated genomic DNA using Q5 Hot Start High-Fidelity DNA Polymerase (M0493L, NEB, Ipswich, MA, USA). PCR products underwent primer walking Sanger sequencing (Sterky and Lundeberg, 2000). For chick, the EcoRI inverse PCR yielded a desired length of 2 kb of promoter sequence but for duck and quail less than 2 kb of the promoter was initially sequenced so inverse PCR was repeated using XbaI (R0145S, NEB, Ipswich, MA, USA).

### Mmp13 Promoter Sequence Analysis

To identify transcription factor binding sites we used the JASPAR 2020 database, which contains transcription factor-binding profiles stored as position frequency matrices (Fornes et al., 2020). To map transcription factor binding sites onto the *Mmp13* promoter sequences of chick, quail, and duck we used TFBSTools (Tan and Lenhard, 2016), which is an R package (Team, 2013). For the proximal region of the *Mmp13* promoter (*i.e.*, −184 bp for chick and quail, and −181 bp for duck) all vertebrate transcription factors were included in the analysis. For the −2 kb promoter region only RUNX2 (ID = MA0511), SMAD3_1 (ID = PB0060), SMAD3_2 (ID = PB0164), SMAD2-SMAD3-SMAD4 (ID = MA0513), SMAD3 (ID = MA0795), SMAD4 (ID = MA1153), and SMAD2/3 (ID = MA1622) were included in the analysis since these are TGFβ activated (Heldin et al., 1997; Derynck et al., 1998; Derynck et al., 2008). Position frequency matrices were converted to position weighted matrices by setting pseudocounts to 0.8 (Nishida et al., 2009), and background frequencies of nucleotides to 0.25. The minimum threshold score was set to 95% for the −184/181 bp promoter region and 90% for the −2 kb promoter region.

A SMAD binding element (*i.e*., 5’-GGC(GC/CG)-3’), which was not annotated in the JASPAR 2020 database, was manually added (Martin-Malpartida et al., 2017). We also identified another possible SMAD binding motif nested entirely within an AP1 binding element in the proximal region of the *Mmp13* promoter of all three species that we excluded from our experimental design and analysis. Sequence logos were generated using the seqLogo function in TFBSTools (Supplemental Figure S1).

### Generation of Mmp13 *and* Runx2 Constructs

Each *Mmp13* promoter sequence was amplified by PCR using Q5 Hot Start High-Fidelity DNA Polymerase (M0493L, NEB, Ipswich, MA, USA). To generate luciferase constructs, pGL3 was digested with HindIII-HF (R3104S, NEB, Ipswich, MA, USA) and XhoI (R0146S, NEB, Ipswich, MA, USA). The amplified *Mmp13* promoter sequences and digested pGL3 were purified using GeneJET PCR Purification Kit and cloned using NEBuilder HiFi DNA Assembly Master Mix (E2621L, NEB, Ipswich, MA, USA). Mutations in *Mmp13* promoter SNPs were generated through site-specific mutagenesis PCR with the mutations in the primers (Ho et al., 1989). All constructs were verified by sequencing and midi-prepped for transfection using PureLink Fast Low-Endotoxin Midi Kit (A36227, Invitrogen, Carlsbad, CA, USA).

To generate *Runx2* overexpression constructs, full length cDNA was synthesized using Maxima H Minus first strand cDNA synthesis kit (K1651, Thermo Fisher Scientific, Waltham, MA, USA) following the manufacturer’s protocol with 2 μg of total HH37 chick, quail, or duck lower jaw RNA and 100 pmol of d(T)20 VN primer (Chu et al., 2020). The cDNA synthesis reaction was carried out at 50°C for 30 min, 55°C for 10 min, 60°C for 10 min, 65°C for 10 min, and 85°C for 5 min. Full length *Runx2* was amplified by PCR using Q5 Hot Start High-Fidelity DNA Polymerase and cloned using CloneJET PCR Cloning Kit. Full length *Runx2* was confirmed by Sanger sequencing and cloned into our pPIDNB custom-made plasmid (Chu et al., 2020), which was digested with AflII (R0520S, NEB, Ipswich, MA, USA) and PstI (R3140S, NEB, Ipswich, MA, USA), using NEBuilder HiFi DNA Assembly Master Mix. The pPIDNB plasmid contains a constitutively active mNeongreen (GFP) (Shaner et al., 2013), which serves as a reporter for transfection or electroporation efficiency; and a doxycyline (dox)-inducible (Gossen et al., 1995; Loew et al., 2010; Heinz et al., 2011) mScarlet-I (RFP) (Bindels et al., 2017). All constructs were verified by sequencing and midi-prepped for transfection or electroporation using PureLink Fast Low-Endotoxin Midi Kit (A35892, ThermoFisher Scientific, Waltham, MA, USA).

### Transfection and Luciferase Assay

Cells were plated at 65,000 cells/cm^2^ in 24-well plates (353047, Corning, Corning, NY, USA). Cells were transfected in each well using 1.5 μl Lipofectamine 3000 (L3000008, Invitrogen, Carlsbad, CA, USA), 1.5 µl P3000 reagent, 150 ng of β-Gal transfection efficiency control construct, and 1500 ng of *Mmp13* promoter luciferase construct, or 750 ng of *Mmp13* promoter luciferase construct when combined with 400 ng of pPIDNB*-Runx2* overexpression construct. Cells were transfected for 18 hours, recovered in complete media conditions for 8 hours, and then serum deprived in DMEM for DF-1 cells and MEM α for CCL-141 cells without FBS for 18 hours. Cells transfected with pPIDNB*-Runx2* were treated with a final concentration of 100 ng/ml of doxycycline hyclate (dox) (446060250, Acros Organics, Fair Lawn, NJ, USA) in DMEM or MEMα without FBS for 24 hours. Cells were lysed in 1X lysis buffer (E1531, Promega, Madison, WI, USA) and analyzed for luciferase activity using beetle luciferin (E1602, Promega, Madison, WI, USA) and Coenzyme A (J13787MF, ThermoFisher Scientific, Waltham, MA, USA) normalized to β-galactosidase activity using Galacto-Star β-Galactosidase Reporter (T1012, Invitrogen, Carlsbad, CA, USA) as previous described (Chen et al., 2012b). Luminescence was measured using a Spectramax M5 luminometer. At least two preparations of each DNA construct were tested for the over-expression experiments.

### Statistics

Statistical analyses and graphing of data were performed using Prism (GraphPad Software, Version 9.0.0). Data are represented as a mean ± standard error of the mean (SEM). Statistical significance was determined through two-tailed ANOVA adjusted for multiple comparisons using the Bonferroni method for the majority of experimental data, or a paired Student’s t-test for comparisons between control and treatment groups for TRAP quantification. For *in ovo* data, n refers to the total number of embryos analyzed per group. For *in vitro* data, n refers to the total number of individual wells analyzed per group, with each experiment replicated at least three times. For qPCR and luciferase experiments, each sample was run in technical duplicate and averaged. If outliers were found within data sets, a standard Q-test was performed with no more than 1 outlier removed from any group. In all figures, p < 0.05 was considered statistically significant, although some statistical comparisons reached significance below p < 0.01, p < 0.001, or p < 0.0001 as noted. Group size “n” is denoted in the figure legends. Formal power analyses were not conducted. Figures were assembled in Adobe Illustrator 2020 (Version 24.2.3).

## RESULTS

### Bone resorption and MMP13 levels are species-specific and spatially regulated

To identify coincident areas of bone resorption and MMP13 localization, we performed TRAP staining and IHC for MMP13 on sections of HH40 quail and duck lower jaws. Previously we have shown that by HH40, species-specific differences in domains of bone resorption are evident in quail and duck tissues (Ealba et al., 2015). We also performed trichrome staining on adjacent sections to label areas of bone deposition (Figure 1A-B). We observe qualitatively higher levels of TRAP staining in quail lower jaws in all bone regions compared to stage-matched duck lower jaws (Figure 1C-E). In the quail angular bone, MMP13 protein is elevated compared to similar regions in duck and overlaps directly with TRAP staining (Figure 1F-K). In the quail dentary bone, areas of TRAP staining overlap with domains of MMP13 and are elevated compared to similar regions in duck, however the duck dentary has more elevated levels of TRAP and MMP13 than in the angular bone (Figure 1L-Q).

**Figure 1.**
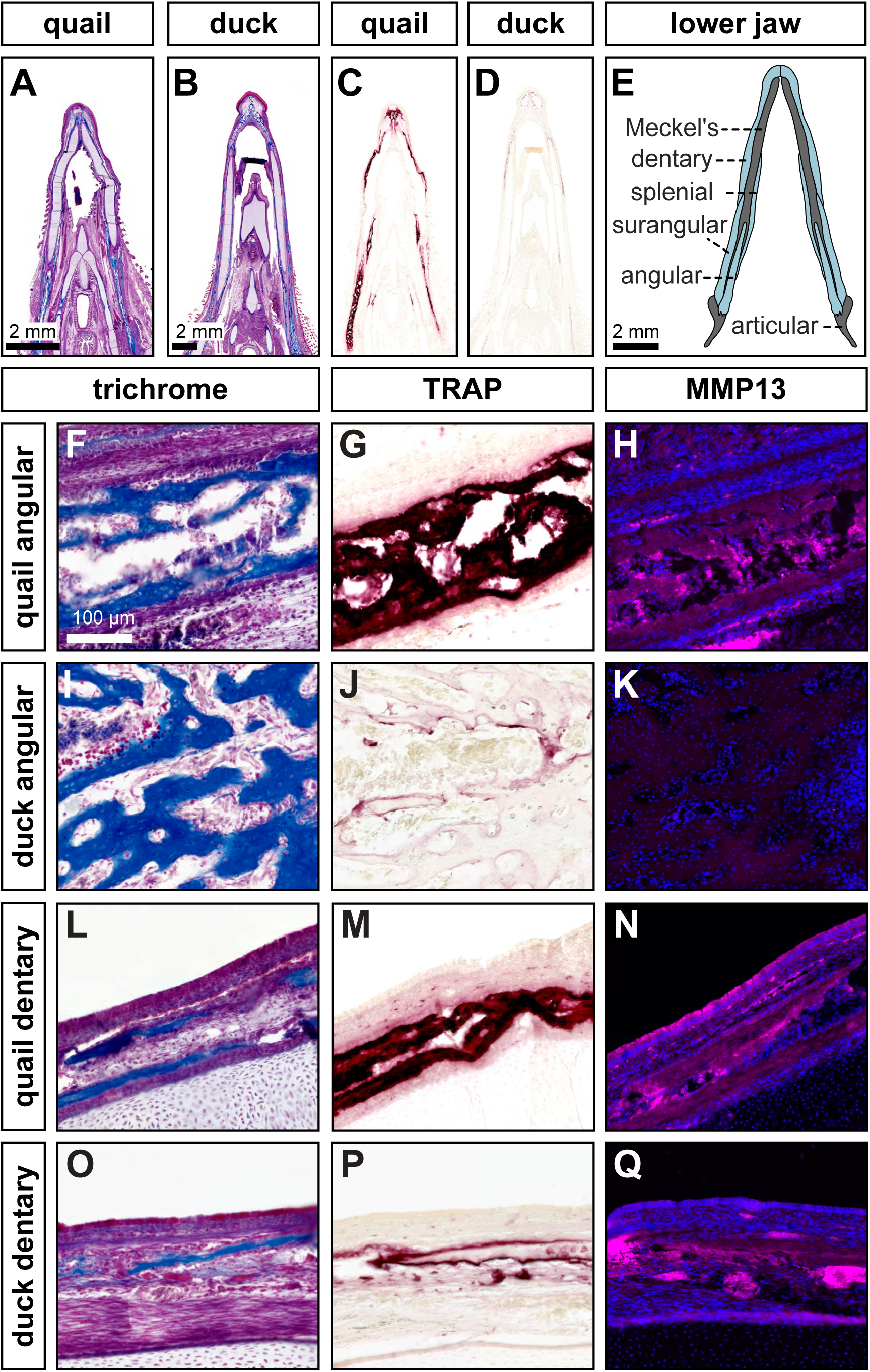
Species-specific differences in spatial domains and levels of TRAP activity and of MMP13 expression. Sections of lower jaws stained with Trichrome (osteoid matrix of bone is blue) in **(A)** quail (n = 4) versus **(B)** duck (n = 4) at HH40. **(C)** Adjacent sections stained for TRAP (red) reveals more robust bone resorption in quail versus **(D**) duck. **(E)** Schematic of bones within the avian lower jaw. **(F)** Adjacent sections through the more proximal angular bone stained with Trichrome, **(G)** TRAP, and **(H)** MMP13 antibody (pink) and Hoechst dye (blue) in quail. **(I)** Angular bone stained with Trichrome, **(J)** TRAP, and **(K)** MMP13 antibody reveal substantially less bone resorption in duck. (**L)** Adjacent sections through the more distal dentary bone stained with Trichrome, **(M)** TRAP, and **(N)** MMP13 antibody in quail. **(O)** Dentary bone stained with Trichrome, **(P)** TRAP, and **(Q)** MMP13 antibody reveals more bone resorption in the dentary versus the angular bone of duck but still less bone resorption overall compared to that observed in quail.

### TGFβ signaling is upregulated in quail during key stages of bone resorption

To examine whether higher levels of MMP13 protein expression and bone resorption observed in quail versus duck correlate with differential regulation of TGFβ signaling, we quantified gene expression of TGFβ ligands, receptors, effectors, and target genes relative to HH31. When examining expression of TGFβ ligands, we find an increase in *Tgfβ1* for chick (2.2-fold, p < 0.05) and quail (2-fold, p < 0.05) and in *Tgfβ3* for chick (2.8-fold, p < 0.0006) and quail (4.8-fold, p < 0.0001) at HH37, which is a stage when bone resorption can be first detected in the lower jaw by TRAP (Ealba et al., 2015). In contrast, we observe no change in duck (Figure 2A-B). *Tgfβ2* levels do not change in chick or quail between HH31 and HH40 but decrease in duck at HH37 (Supplemental Figure S2A; 3.4-fold, p < 0.05). For TGFβ receptors, *Tgfβr1, Tgfβr2,* and *Tgfβr3* mRNA expression does not change over time in any species, whereas the non-canonical receptor *Acvrl1* increases at HH37 in quail with no changes in chick or duck from HH31 to HH40 (Figure 2C, Supplemental Figure S2B-D). For downstream effectors, *Smad2* increases at HH37 in chick (2.5-fold, p < 0.0001) and quail (2.8-fold, p < 0.05), but decreases in duck (1.6-fold, p < 0.05), whereas *Smad3* decreases at HH40 in all species (Figure 2D, Supplemental Figure S2E;). We find that when comparing gene expression at HH37, quail has higher expression of *Tgfβ1 (*5.6-fold, p < 0.0004*), Tgfβ3* (3.4-fold, p < 0.009), *Tgfβr1* (6.5-fold, p < 0.001), *Acvrl1* (3.5-fold, p < 0.0001), and *Smad2* (3-fold, p < 0.0001) compared to duck. No differences are found between quail and duck in *Tgfβ2*, *Tgfβr2*, *Tgfβr3*, and *Smad3*. Chick levels of TGFβ signaling show similar trends to those in quail, however we do observe significant differences in *Tgfβ1* (4-fold, p < 0.0004)*, Tgfβ3* (1.7-fold, p < 0.0007), *Tgfβr1* (1.8-fold, p < 0.0003), *Acvrl1* (5-fold, p < 0.0001), *Smad2* (2-fold, p < 0.0002), and *Smad3* (3.3-fold, p < 0.003) compared to quail. When comparing duck to chick, we observe significant differences in *Tgfβ3* (2-fold, p < 0.01), *Tgfβr1* (3.7-fold, p < 0.001), *Smad2* (6.8-fold, p < 0.0001), and *Smad3* (2.9-fold, p < 0.004).

**Figure 2.**
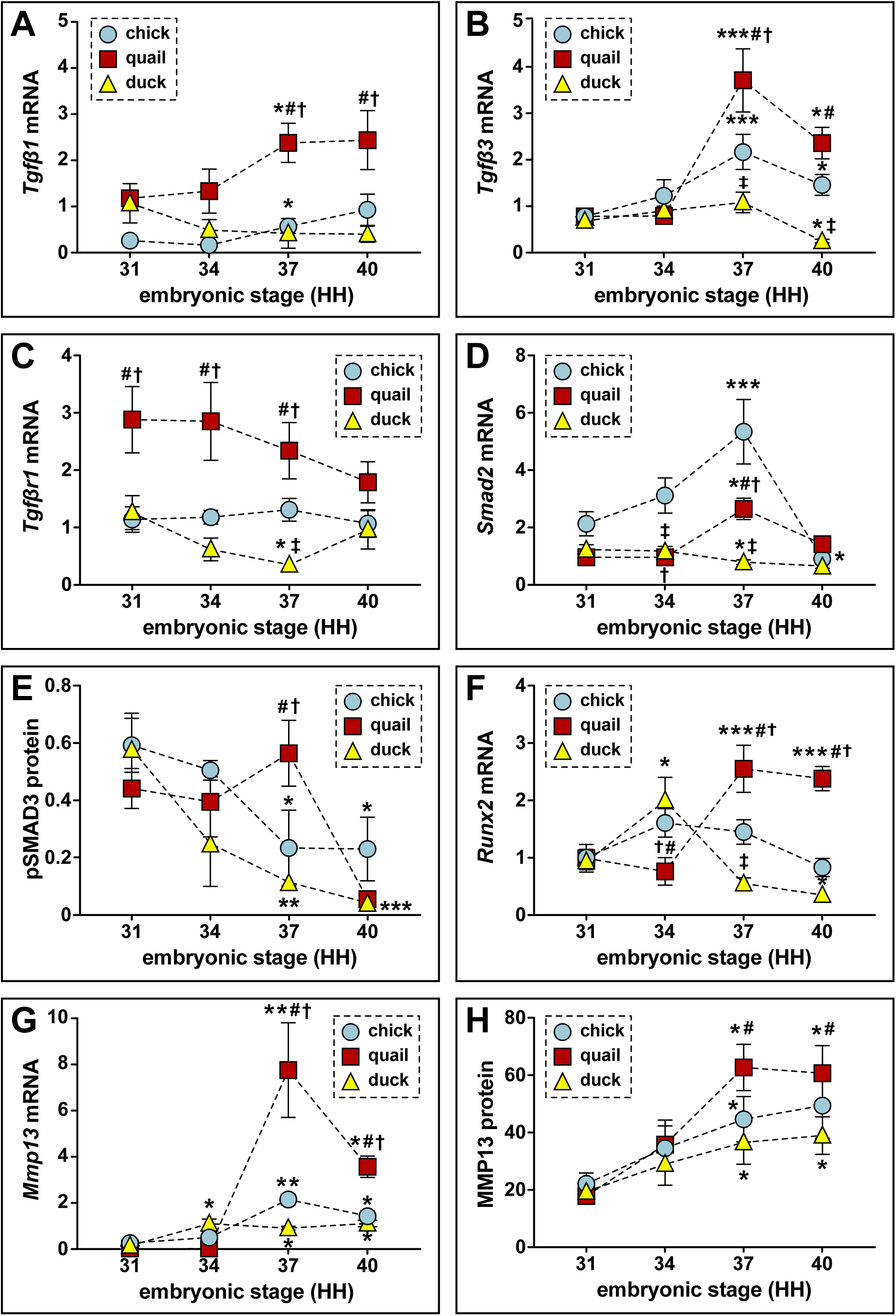
Relative mRNA and protein levels of TGFβ pathway members and targets in the developing lower jaws of chick, quail, and duck. **(A)** RT-qPCR analyses show that *Tgfβ1* mRNA increases 1.5-fold in chick (blue circles, n = 8) and 2-fold in quail (red squares, n = 8) at HH37, and does not change in duck (yellow triangles, n = 10) in later developmental stages; *Tgfβ1* is higher in quail than chick or duck at HH37. (B) *Tgfβ3* mRNA increases 2.8-fold in chick and 4.8-fold in quail at HH37, while duck levels decrease by HH40. Chick and quail have higher *Tgfβ3* at HH37 compared to duck. (C) *Tgfβr1* mRNA remains constant for chick and quail whereas duck decreases significantly at HH37. Quail maintain higher levels of *Tgfβr1* from HH31 to HH37 relative to chick and duck. (D) *Smad2* mRNA increases 2.5-fold in chick and 2.8-fold in quail at HH37, whereas duck levels decrease 1.6-fold. At HH40, chick and quail robustly decrease compared to HH37. **(E)** Western blots show pSMAD3 protein levels are elevated 5-fold in quail (n = 12) at HH37. Duck (n = 12) levels decrease over time, while chick (n = 9) levels significantly decrease at HH37 and HH40 compared to HH31. Quail has higher levels at HH37 than duck. (F) *Runx2* mRNA increases 2.6-fold in quail at HH37 and remains elevated at HH40, whereas duck increases at HH34 and then trends downward from HH37 to HH40. Chick *Runx2* does not change. (G) *Mmp13* mRNA increases 8.4-fold in chick and 1300-fold in quail at HH37, whereas duck increases 8-fold at HH34 and remains elevated until HH40. **(H)** MMP13 protein increases 2-fold in chick, 3-fold in quail, and 1.8-fold in duck at HH37 and remains elevated at HH40. p < 0.05 and * denotes significance from HH31 within each group, # denotes significance between quail and duck at same stage, † denotes significance between chick and quail, ‡ denotes significance between chick and duck.

To confirm activation of the TGFβ pathway, we assayed for phosphorylated (p) SMAD3 and observe a 5-fold (p < 0.004) higher level in quail versus duck at HH37, indicating that quail have elevated TGFβ signaling at this stage (Figure 2E, Supplemental Figure S3A). Correspondingly, we find significant upregulation of TGFβ targets including *Runx2* (4.6-fold, p < 0.0001), *Mmp13* (8.5-fold, p < 0.0001), *Pai1* (5.3-fold, p < 0.0001), and *Mmp2* (9.3-fold, p < 0.0001) in mRNA in quail versus duck at HH37, as well as MMP13 (1.6-fold, p < 0.01) protein levels (Figure 2F-H, Supplemental Figure S3F-G, Supplemental Figure S3B). Gene expression increases between HH34 and HH37 for *Runx2*, *Mmp13*, *Pai1*, and *Mmp2* in quail, which mirrors the increases in TGFβ ligand expression and higher activation of pSMAD3, whereas in duck these genes either decrease or remain flat during the same transition. Chick gene expression follows similar trends to that observed in quail, with higher expression of *Runx2* (2.6-fold, p < 0.01) but no difference with *Mmp13* or *Pai1* compared to duck. We also observe an increase in the osteoclast-secreted *Mmp9* in all species at HH37, with a 2.3-fold higher expression in quail compared to duck (p < 0.001), and a 1.5-fold higher expression in quail than chick (Supplemental Figure S2H; p < 0.01). *Mmp14*, which is mostly secreted by osteocytes (Wu et al., 2009; Qing et al., 2012; Dole et al., 2017), does not change in expression and is relatively similar among all species, suggesting differential regulation of *Mmps* (Supplemental Figure S4A). *Ctsk*, which is secreted by osteoclasts and osteocytes (Qing and Bonewald, 2009; Qing et al., 2012; Dole et al., 2017; Lotinun et al., 2019), follows a similar trend as other bone resorption markers with increases at HH37 in chick (3.2-fold, p < 0.05) and quail (1.5-fold, p < 0.05, but not duck (Supplemental Figure S4B). *Sost*, a marker for osteocytes (Zhu et al., 2011; Hernandez et al., 2014), increases at HH40 in chick (7.6-fold, p < 0.01) with no changes in quail or duck from HH34 to HH40 (Supplemental Figure S4C). Taken together, we observe that members of the TGFβ signaling pathway are more highly expressed and show greater activation in chick and quail versus duck.

### Sensitivity to TGFβ signaling is cell autonomous and species-specific

To test if the greater activation of TGFβ pathway members and targets observed in chick and quail versus duck lower jaws is due to intrinsic species-specific differences in sensitivity to TGFβ signaling, we performed experiments comparing the response of chick and duck embryonic fibroblasts to rTGFβ1. We treated a chick cell line (*i.e.*, DF-1) and a duck cell line (*i.e.,* CCL-141) with rTGFβ1 and assayed for activation of target genes. We observe a 3-fold (p < 0.001) induction in pSMAD3 protein levels in chick cells treated with rTGFβ1 compared to controls but only observe a 1.7-fold (p < 0.07) increase, which is not significant in duck cells relative to controls (Figure 3A, Supplemental Figure S3C). We also observe a species-specific response for *Runx2* and *Mmp13* (Figure 3B-C). In chick cells, *Runx2* increases 3-fold at 3-hours (p < 0.02) and 6-hours (p < 0.0005) post-treatment and shows a 7.4-fold induction (p < 0.0001) at 24 hours. However, in duck cells, we find no significant response until 24 hours when we observe a 1.8-fold induction (p < 0.04), which is much less than the induction in chick (p < 0.0001). We observe a similar response in chick cells for *Mmp13* with a 4.2-fold induction (p < 0.0004) at 6 hours and a 10.8-fold induction (p < 0.0001) at 24 hours. In contrast, we only observe a 3.7-fold induction (p < 0.008) of *Mmp13* expression in duck cells at 24 hours, which is less than the 10-fold induction at this time point in chick cells (p < 0.0001). MMP13 protein levels parallel the gene expression response with a 1.6-fold induction (p < 0.001) in chick, but no induction in duck cells (Figure 3D, Supplemental Figure S3D). We do not observe a similar effect when we examine the response of other targets such as *Pai1* and *Mmp2*. For duck cells treated with rTGFβ1, *Pai1* increases 2-fold (p < 0.04) at 1 hour and 2.4-fold (p < 0.02) at 24 hours, whereas in chick *Pai1* does not increase until 24 hours with a 3.5-fold induction (p < 0.0001; Supplemental Figure S5A). *Mmp2* increases 3.5-fold (p < 0.05) in duck cells at 1 hour and 7.5-fold (p < 0.0001) at 24 hours post-treatment in duck cells, whereas chick has a later and dampened response with a 2-fold induction at 6-hours (p < 0.05) and 24-hours (p < 0.05) post-treatment (Supplemental Figure S5B).

**Figure 3.**
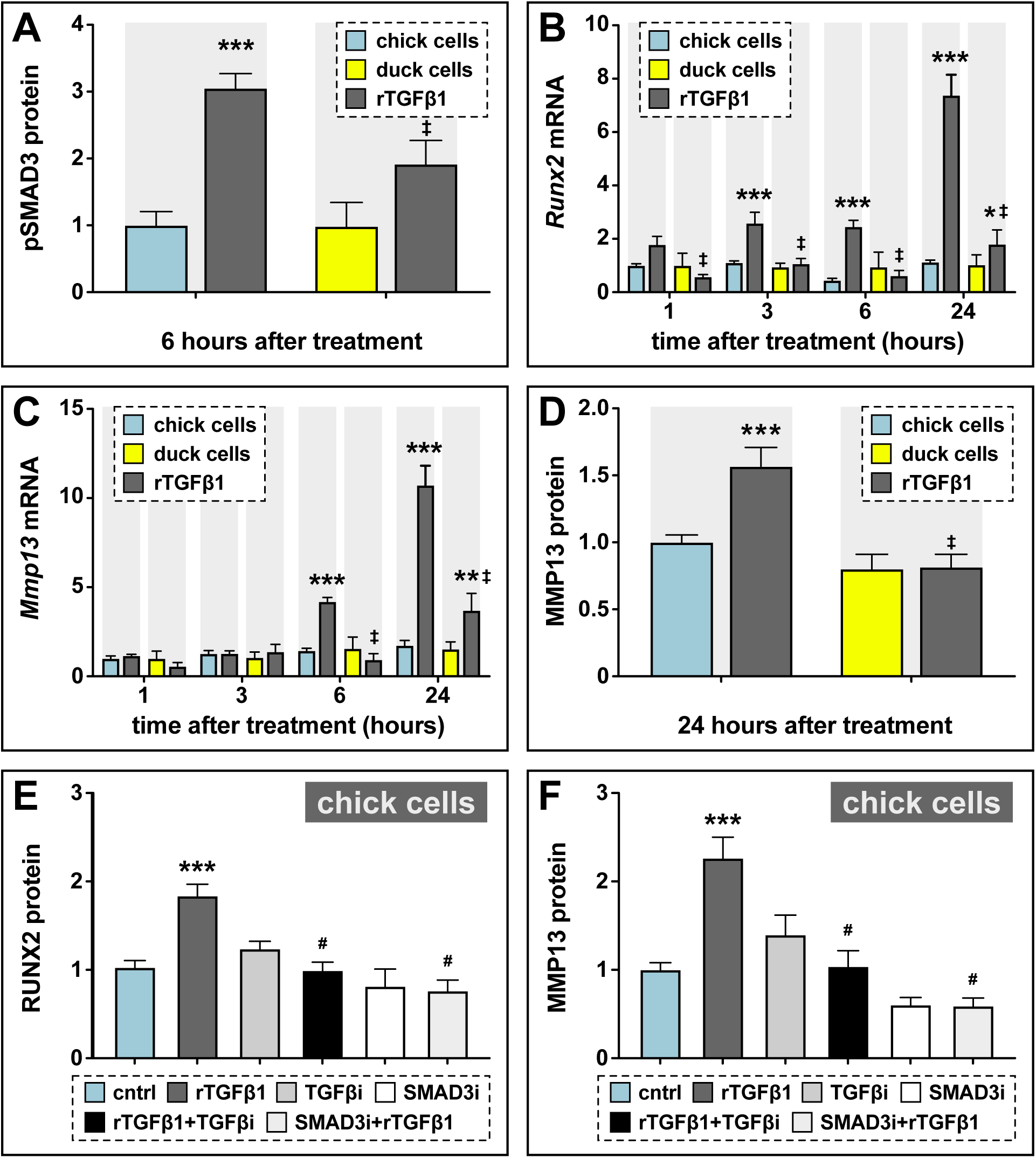
Sensitivity of chick and duck cells to rTGFβ1 and effects of TGFβR1 and SMAD3 inhibition on RUNX2 and MMP13. **(A)** pSMAD3 protein levels in cells treated for 2 hours with 5 ng/ml rTGFβ1 (dark gray) in chick (blue) and duck (yellow) cells show a significant induction in chick (n = 8). **(B)** Chick and duck cells treated with rTGFβ1 for 1 to 24 hours. Chick *Runx2* mRNA increases 3-fold with rTGFβ1 treatment at 3 and 6 hours and 8-fold at 24 hours, while duck *Runx2* does not increase until 24 hours. In rTGFβ1 treated cells, duck have significantly lower *Runx2* at every time point compared to chick (n = 8). **(C)** Chick *Mmp13* mRNA increases 4.5-fold with rTGFβ1 treatment at 6 hours and 10-fold at 24 hours, while duck *Runx2* does not increase until 24 hours. In rTGFβ1 treated cells, duck has lower *Mmp13* at every time point compared to chick (n = 8). **(D)** MMP13 protein in cells treated for 24 hours with rTGFβ1 show a 1.5-fold induction in chick but no response in duck (n = 8). **(E)** RUNX2 protein levels in chick cells treated for 24 hours with rTGFβ1 (dark gray), TGFβR1 inhibitor (medium gray), a combination of both rTGFβ1 and TGFβR1 inhibitor (black), SMAD3 inhibitor (white), and a combination of both rTGFβ1 and SMAD3 inhibitor (light gray). RUNX2 protein increases 2-fold with rTGFβ1 but when rTGFβ1 is combined with either a TGFβr1 or a SMAD3 inhibitor, there is a significant decrease compared to rTGFβ1 alone (n = 12). **(F)** MMP13 protein increases 2.3-fold with rTGFβ1 treatment but when rTGFβ1 is combined with either a TGFβR1 or a SMAD3 inhibitor, there is a significant decrease compared to rTGFβ1 alone (n = 12). * denotes significance from control p < 0.05, ** denotes significance from control p < 0.01, *** denotes significance from control p < 0.001, # denotes significance from rTGFβ1, † denotes significance between chick and quail, ‡ denotes significance between chick and duck.

To assess if the observed induction of *Runx2* and *Mmp13* mRNA by rTGFβ1 is mediated via the canonical TGFβ signaling pathway, we utilized small molecule inhibitors of TGFβR1 (*i.e.*, SB431542) and SMAD3 (*i.e.,* SIS3). We treated chick cells with rTGFβ1 and/or SB431542, or rTGFβ1 and/or SIS3 for 24 hours and measured the response on *Runx2* and *Mmp13* protein levels. rTGFβ1 treatment induces a 2-fold increase in RUNX2 (p < 0.0001) and MMP13 (p < 0.0001), whereas inhibitor treatment alone has no effect (Figure 3E-3F, Supplemental Figure S3E-F). However, the ability of rTGFβ1 to induce RUNX2 and MMP13 is abolished when cells are treated with rTGFβ1 in combination with either inhibitors of TGFβR1 or SMAD3 (Figure 3E-F). We observe no effect on MMP2 protein levels in cells treated with inhibitors of TGFβR1 or SMAD3 (data not shown).

To test if the species-specific differences in sensitivity of chick and duck cells to TGFβ signaling also occur at the tissue level and in the context of development, we dissected HH34 lower jaws from chick, quail, and duck (Figure 4A-C) and treated them with a series of rTGFβ1 concentrations. We observe no induction of TGFβ pathway members with 5 ng/ml rTGFβ1 treatment in the lower jaws of chick, quail, and duck (data not shown), but observe species-specific differences in the response of TGFβ pathway members to concentrations of 10, 25, and 50 ng/ml. We find elevated levels of pSMAD3 in chick (2.3-fold, p < 0.01) and quail (1.8-fold, p < 0.05) versus duck with 25 ng/ml rTGFβ1 treatment for 6 hours (Figure 4D, Supplemental Figure S3G). For 50 ng/ml, we only observe an induction in quail (3.4-fold, p < 0.002), with a trending increase in chick (3-fold, p < 0.059), but no response in duck lower jaws. Quail pSMAD3 levels are higher compared to duck for 50 ng/ml (2.3-fold, p < 0.02). *Runx2* and *Mmp13* mRNA expression increase in chick (2.4-fold, p < 0.03; and 2.5-fold, p < 0.0001) and quail (3.2-fold, p < 0.0001; and 3.2-fold, p < 0.0001), but not in duck following treatments with 25 ng/ml rTGFβ1 for 24 hours (Figure 4E-F). Treatment with rTGFβ1 increases *Runx2* in chick (3.2-fold, p < 0.0001) and quail (4,3-fold, p < 0.0001), and *Mmp13* in chick (5-fold, p < 0.0001) and quail (6.4-fold, p < 0.0001) relative to duck. MMP13 protein levels only show increases in quail at 25 ng/ml, but no induction is observed in chick or duck at any dose (Figure 4G, Supplemental Figure S3H). To test if rTGFβ1 treatment affects TGFβ receptor expression, we assayed for changes in expression of *Tgfβr1*, *Tgfβr2,* and *Tgfβr3.* We find no change in expression for these receptors in any species (Supplemental Figure S5C-D and data not shown). To test if rTGFβ1 treatment stimulates other known TGFβ target genes, we examined *Pai1* and *Mmp2*. *Pai1* mRNA expression in chick and duck does not change with rTGFβ1 treatment, whereas we observe a 1.8-fold induction (p < 0.003) in quail (Supplemental Figure S5E). However, *Pai1* expression is higher in chick (2.1-fold, p < 0.0001) and quail (2.4-fold, p < 0.0001) compared to duck when treated with rTGFβ1. *Mmp2* is induced in chick (1.5-fold, p < 0.02) and quail (1.5-fold, p < 0.04) but not in duck lower jaws (Supplemental Figure S5F). To examine if treatment with rTGFβ1 induces markers for bone formation in lower jaws within 24 hours, we assayed for changes in *Col1a1* and *Ocn*. We do not observe changes in *Col1a1* levels with rTGFβ1 treatment in any species, however, we do observe a 1.8-fold decrease (p < 0.05) in *Ocn* levels in duck (Supplemental Figure S5G-H).

**Figure 4.**
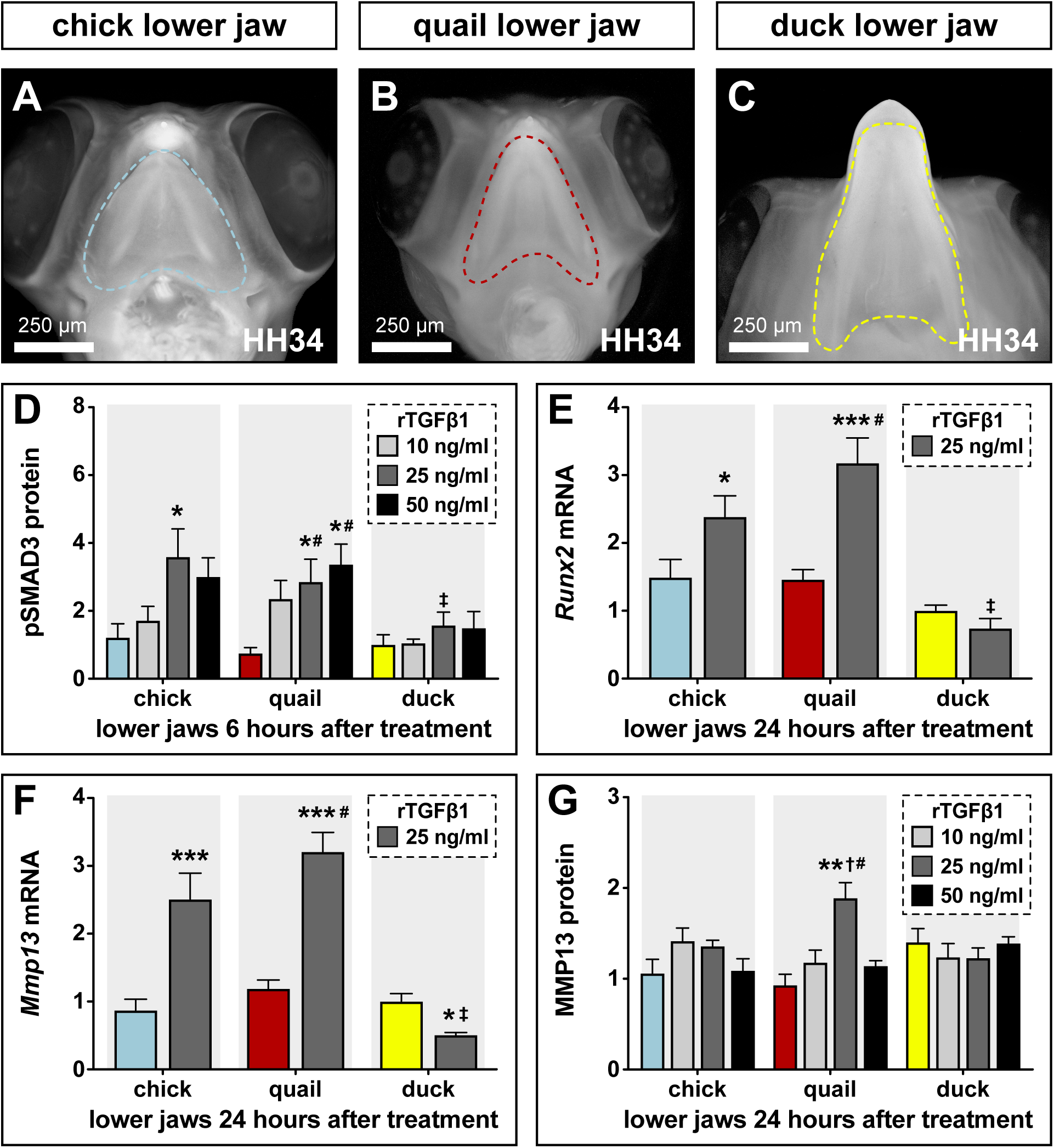
Sensitivity of chick, quail, and duck lower jaws to rTGFβ1 and the effects on RUNX2 and MMP13. **(A)** HH34 chick, **(B)** quail, and **(C)** duck heads in ventral view showing the boundaries (dashed line) of the lower jaws dissected for analyses and culture experiments. **(D)** Chick (blue), quail (red), and duck (yellow) lower jaws treated in culture with 10 (medium gray), 25 (dark gray), and 50 (black) ng/ml of rTGFβ1 for 6 hours. pSMAD3 levels increase 3.6-fold in chick and 3-fold in quail at 25 ng/ml, while duck shows no induction at any time point following treatment. Quail pSMAD3 levels are significantly induced with 50 ng/ml treatment (n = 12). **(E)** Chick, quail, and duck lower jaws treated with 25 ng/ml rTGFβ1 for 24 hours. *Runx2* mRNA increases 2.4-fold in chick and 3.2-fold in quail, whereas duck significantly decreases by 1.3-fold (n = 10). **(F)** Chick, quail, and duck lower jaws treated with 25 ng/ml rTGFβ1 for 24 hours. *Mmp13* mRNA increases 2.5-fold in chick and 3.2-fold in quail, whereas duck significantly decreases by 2-fold (n = 10). **(G)** Chick, quail, and duck lower jaws treated in culture with 10, 25, and 50 ng/ml of rTGFβ1 for 24 hours. MMP13 protein levels show a 2-fold induction in quail with 25 ng/ml rTGFβ1 but no response in chick or duck. * denotes significance from HH31 within each group p < 0.05, ** denotes significance from HH31 within each group p < 0.01, # denotes significance between quail and duck at same stage, † denotes significance between chick and quail, ‡ denotes significance between chick and duck.

### Bone resorption requires TGFβ signaling in the jaw skeleton

To determine the extent to which members and targets of the TGFβ pathway regulate bone resorption, we dissected HH35 lower jaws from quail; treated them on the right side with beads soaked in inhibitors of TGFβR1 (Figure 5A), SMAD3 (Figure 5B), or MMP13 (Figure 5C); cultured them in osteogenic media for 5 days, and then performed TRAP staining. Vehicle control beads were implanted on the left side of each sample.

**Figure 5.**
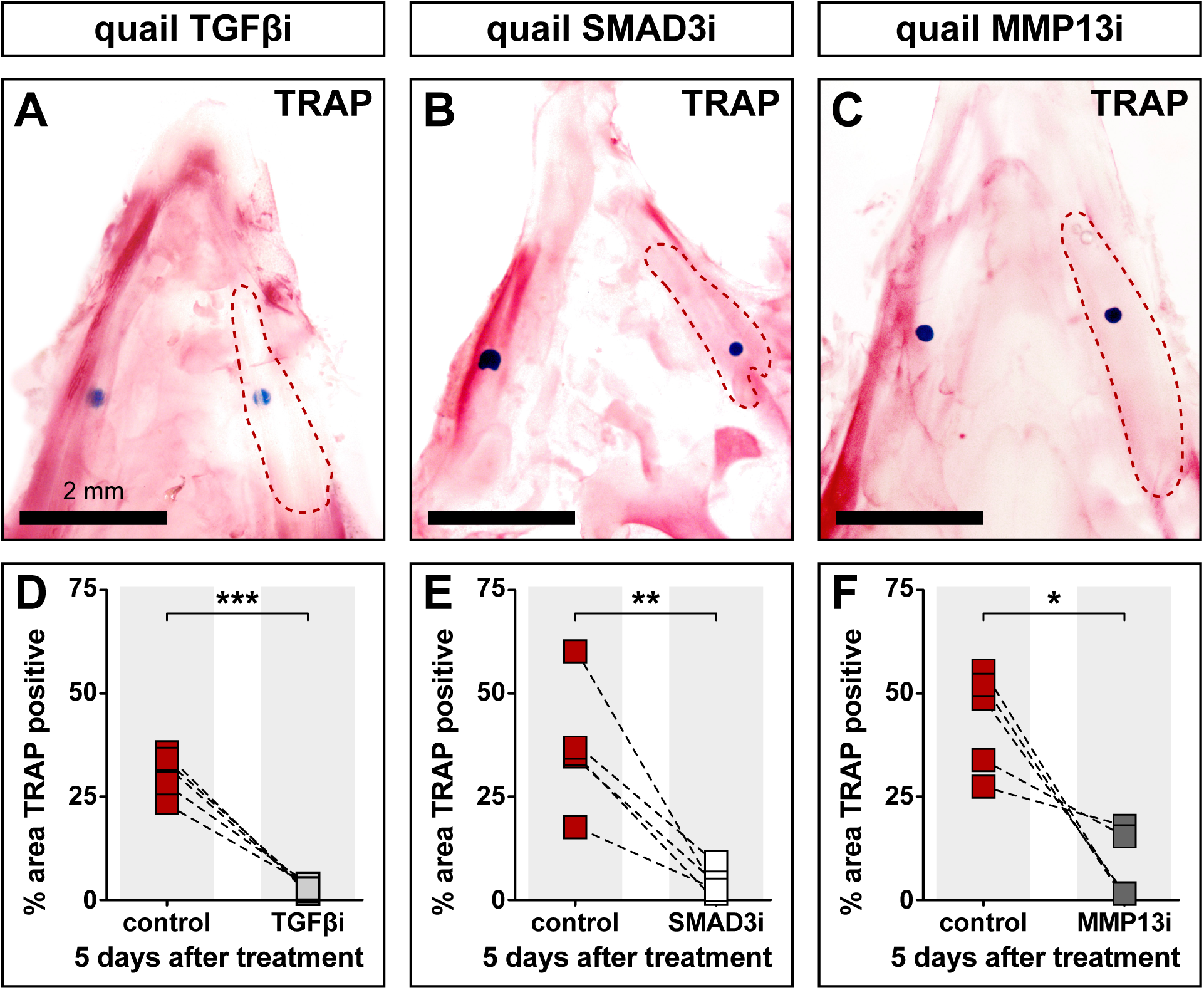
Effects of TGFβR1, SMAD3, and MMP13 inhibition on TRAP staining. Quail lower jaws harvested at HH35 and placed in culture with control beads (left side) and treatment beads (right side) soaked in **(A)** TGFβR1 inhibitor (TGFβi, n = 5), **(B)** SMAD3 inhibitor (SMAD3i, n = 5), and **(C)** MMP13 inhibitor (MMP13i, n = 5). The effects of inhibitor treatments can be seen on TRAP staining (red) after 5 days of culture (red dashed lines). Quantification of the % area of TRAP-positive staining for **(D)** TGFβR1 inhibitor (medium gray), **(E)** SMAD3 inhibitor (white), and **(F)** MMP13 inhibitor (dark gray) in comparison to the contralateral control side. * denotes significance from control within each group p < 0.05, ** denotes significance from control within each group p < 0.01, *** denotes significance from control within each group p < 0.001.

Our results show that the area of TRAP staining around the bead is decreased in quail lower jaws by 28% following TGFβR1 inhibition (p > 0.0004), 33% following SMAD3 inhibition (p > 0.009), and 36% following MMP13 inhibition (p > 0.02) when compared to the area of TRAP staining around the control bead on the contralateral side (Figure 5D-F).

### Species-specific response to TGFβ signaling is mediated by the Mmp13 promoter

To determine the extent to which the *Mmp13* promoter itself may regulate the differential and species-specific response of *Mmp13* to TGFβ signaling, we sequenced a 2 kb fragment immediately upstream from the transcriptional start site of the chick, quail, and duck *Mmp13* promoters (Figure 6A). Chick cells were transfected with either the duck or quail *Mmp13* promoter attached to luciferase. To assess the effects of TGFβ signaling on the regulation of the *Mmp13* promoter, cells were treated with rTGFβ1, a TGFβR1 inhibitor, or a combination of both. For the duck *Mmp13* promoter, we find that treatment with rTGFβ1 has no effect on promoter activity, inhibiting TGFβR1 decreases promoter activity (2.1-fold, p < 0.0001), and a combination of rTGFβ1 and TGFβR1 inhibition decreases promoter activity (Figure 6B; 2.4-fold, p < 0.0001). In contrast, for the quail *Mmp13* promoter, we find a 2-fold activation (p < 0.0001) following treatment with rTGFβ1 but no change in promoter activity with TGFβR1 inhibition. No change in promoter activity is observed with a combination of rTGFβ1 and TGFβR1 inhibition compared to untreated cells but activity is lower compared to rTGFβ1 treated cells (2.2-fold, p < 0.0001). Additionally, to assess the effects of *Runx2* on the regulation of the *Mmp13* promoter, we co-transfected cells with −2 kb of the duck or quail *Mmp13* promoters attached to luciferase, as well as a species-specific *Runx2* overexpression construct (Chu et al., 2020). We find that overexpressing duck *Runx2* has no effect on the duck *Mmp13* promoter, however, overexpressing quail *Runx2* induces the activity of the quail *Mmp13* promoter by 2.8-fold (Figure 6C; p < 0.0001).

**Figure 6.**
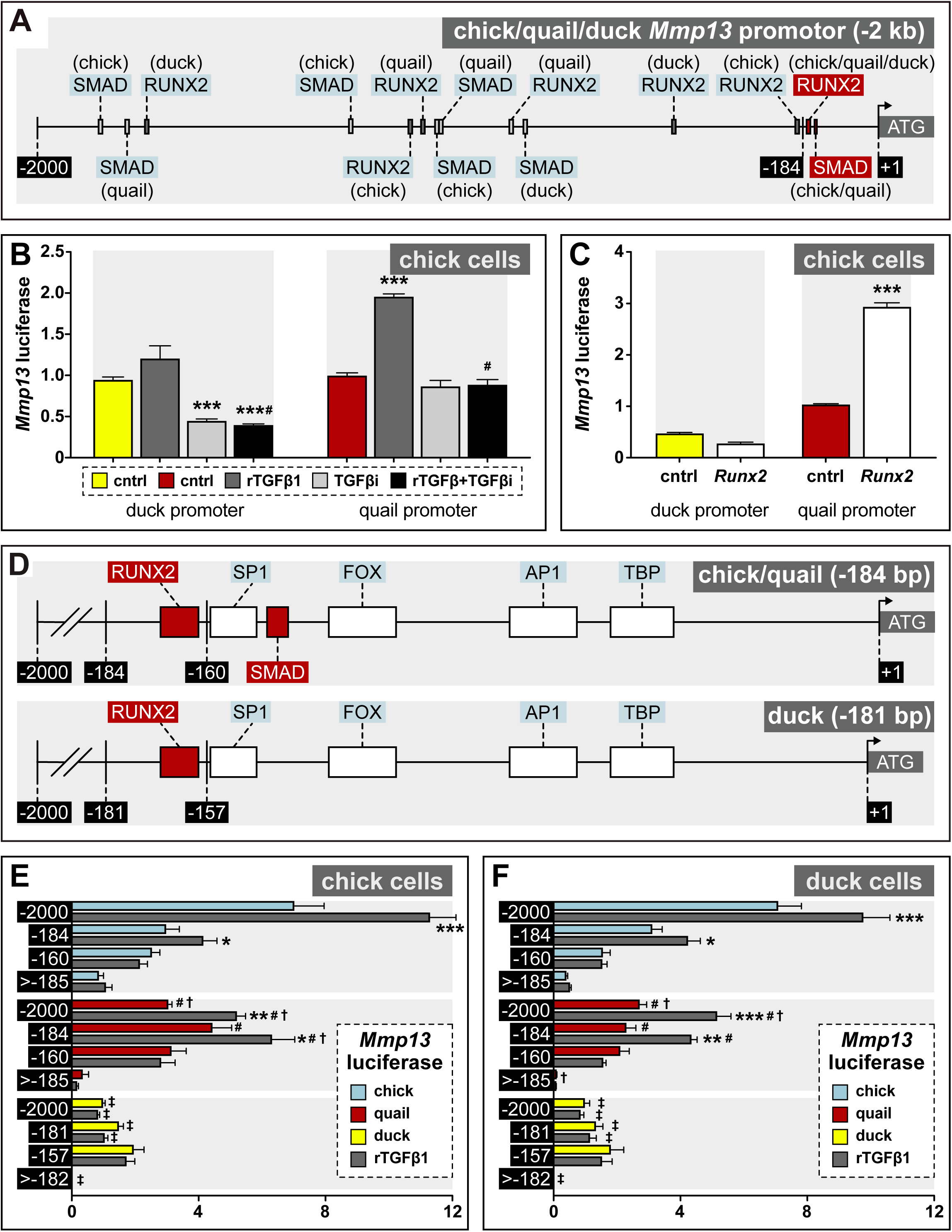
Regulation of *Mmp13* via species-specific promoter elements. **(A)** Schematic of a −2 kb region of the chick, quail, and duck *Mmp13* promoter upstream from the transcriptional start site (ATG). There are four SMAD binding elements in chick, three in quail, and one in duck; and three RUNX2 binding elements in chick, quail, and duck. **(B)** Chick cells transfected with the −2 kb fragment of the duck (yellow) or quail (red) *Mmp13* promoters attached to a luciferase reporter and treated for 24 hours with rTGFβ1 (dark gray), TGFβR1 inhibitor (medium gray), or a combination of both rTGFβ1 and TGFβR1 inhibitor (black). rTGFβ1 treatment has little effect on the duck *Mmp13* promoter but induces activity in the quail promoter. TGFβR1 inhibitor as well as a combination of the TGFβR1 inhibitor and rTGFβ1 decreases activity of the duck *Mmp13* promoter. The inductive effects of rTGFβ1 on the *Mmp13* promoter are abolished in quail when rTGFβ1 is combined with TGFβR1 inhibitor (n = 18). **(C)** Chick cells transfected with duck (yellow) or quail (red) *Mmp13* promoters plus either an empty vector (cntrl) or with a *Runx2*-overexpression plasmid. *Runx2* overexpression has little effect on the duck *Mmp13* promoter but induces quail *Mmp13* promoter activity (n = 18). **(D)** Schematic of the −184/181 b most proximal region of the chick, quail, and duck promoters. Chick, quail, and duck have similar binding elements for Tata Binding Protein (TBP), Activator Protein-1 (AP1), Forkhead Box (FOX), Specificity Protein-1 (SP1), and RUNX2. However, chick and quail contain a SMAD binding element whereas duck does not. **(E)** Chick and **(F)** duck cells transfected with either a −2 kb fragment; a −184 bp fragment including one RUNX2 and one SMAD binding element in chick (blue) and quail; a −181 bp fragment including only a RUNX2 binding element in duck; a −160 bp fragment with no RUNX2 binding elements but including a SMAD binding element in chick and quail; a 157 bp fragment without RUNX2 or SMAD binding elements in duck; a −1815 bp fragment for chick and quail; or a −1818 bp fragment for duck including all binding elements upstream of the first −184/181 bp (*i.e.,* >-185/182). rTGFβ1 treatment (dark gray) induces activity of the −2 kb and −184 bp fragment for chick and quail, however no induction is observed in the duck promotor. In the −160 bp fragment, rTGFβ1-induction is abolished in chick and quail. In the 2000-185 bp fragment, very little activity is observed (n = 24). * denotes significance from untreated control within each group p < 0.05, ** denotes significance from empty vector control within each group p < 0.01, *** denotes significance from empty vector control within each group p < 0.001, # denotes significance between quail and duck, † denotes significance between chick and quail, ‡ denotes significance between chick and duck.

### Differential regulation of Mmp13 is due to species-specific promoter elements

To identify potential regulatory mechanisms underlying these species-specific differences in sensitivity to TGFβ signaling and in levels of *Mmp13* expression, we compared the structure and function of the *Mmp13* promoter in chick, quail, and duck. We analyzed each promoter utilizing the JASPAR 2020 vertebrate transcription factor binding motif data set (Fornes et al., 2020). We mapped potential transcription factor binding elements and uncovered several species-specific differences (Figure 6A). We focused primarily on SMAD and RUNX2 binding elements. Across the −2 kb promoter fragment, we find five SMAD binding elements in chick, four in quail, and three in duck; and three RUNX2 binding elements in chick, quail, and duck. These binding elements are distributed across the *Mmp13* promoter at distinct locations in each species (Figure 6A, Supplemental Table 3). Additionally, one of the SMAD binding elements that we find present in the chick and quail proximal promoter but absent in duck (Figure 6D) is a SMAD binding element not present in the JASPAR 2020 database (*i.e*., 5’-GGC(CG/GC)-3’) previously shown to be regulated by SMAD3 (Martin-Malpartida et al., 2017). We also identified another possible SMAD binding motif nested entirely within an AP1 binding element in the proximal region of the *Mmp13* promoter. We excluded this potential SMAD site from our experimental design and analysis given the functional importance of AP1 binding for the regulation of *Mmp* expression (Benbow and Brinckerhoff, 1997; Pendas et al., 1997; Chakraborti et al., 2003; Selvamurugan et al., 2004b; Samuel et al., 2007; Singh et al., 2010; Hashimoto et al., 2013) and the fact that we could not mutate the shared SMAD motif without simultaneously modifying the AP1 binding element.

To determine the extent to which these SMAD and RUNX2 binding elements account for the differential and species-specific response of *Mmp13* to TGFβ signaling, we generated four sets of differently-sized fragments of the *Mmp13* promoter each driving luciferase expression (Figure 6A and 6D): 1) A −2 kb fragment contains all of the SMAD and RUNX2 binding elements (as described above); 2) a −184 bp fragment contains one RUNX2 and one SMAD binding element in chick and quail but a −181 bp fragment contains only a RUNX2 binding element in duck; 3) a −160 bp fragment has no RUNX2 binding elements but contains a SMAD binding element in chick and quail whereas a - 157 bp fragment contains no RUNX2 or SMAD binding elements in duck; and 4) a - 1815 bp fragment for chick and quail and a −1818 bp fragment for duck contain all the binding elements upstream of the first −184/181 bp (*i.e.,* >-185/182 bp) but without the minimal promoter. We transfected each promoter fragment into chick and duck cells, treated them with rTGFβ1 for 24 hours, and measured luciferase activity.

We observe the highest endogenous activity and induction by rTGFβ1 with the −2 kb promoters of chick (1.6-fold, ±p < 0.0001) and quail (1.7-fold, p < 0.0004), whereas the duck −2 kb promoter shows low activity with no response to rTGFβ1 in chick cells (Figure 6E). In duck cells (Figure 6F), we observe a similar response for the −2 kb promoter of chick (1.4-fold, p < 0.0001) and quail (1.7-fold, p < 0.0004). For the −184 bp chick promoter, endogenous activity is approximately half of that observed with the −2 kb chick promoter. However, we observe a response to rTGFβ1 for the −184 bp promoter of chick (1.4-fold, p < 0.05) and quail (1.4-fold, p < 0.02) in chick cells, and for the −184 bp promoter of chick (1.4-fold, p < 0.05) and quail (1.9-fold, p < 0.0001) in duck cells. In contrast, there is no response to rTGFβ1 observed for the duck −181 bp promoter in either chick or duck cells. For the −160/157 bp promoters, which remove all RUNX2 binding elements from all species, we observe reduced activity for quail and chick but a trending increase in activity for duck. The fragments lacking RUNX2 binding elements are not sensitive to rTGFβ1. When we examined the −1815/1818 bp fragments, which lie upstream of the first −184/181 bp (*i.e.,* >-185/182 bp) without the minimal promoter, we observe reduced or no activity across all species and no response to rTGFβ1. Overall, we find the −2 kb and −184 bp promoter fragments from chick and quail have the highest activity, and are strongly induced by rTGFβ1, whereas the duck promoter shows lower activity and is not responsive to treatments with rTGFβ1. Moreover, the loss of a RUNX2 binding element leads to an increase in *Mmp13* promoter activity for duck.

### SNPs by a RUNX2 binding element affect the species-specific activity of Mmp13

Our comparative analyses of the *Mmp13* promoter identified two SNPs that distinguish chick and quail (*i.e*., “AG”) from duck (*i.e*., “CA”) directly adjacent to a RUNX2 binding element within the −184/181 fragments (Figure 7A). To test if these SNPs affect the differential and species-specific response of the *Mmp13* promoter, we put the chick/quail “AG” SNPs into the duck promoter and the duck “CA” SNPs into the chick/quail promoter. In a second set of constructs we added the SMAD binding element to the duck *Mmp13* promoter and replaced the SMAD binding element in the chick/quail promoter with duck sequence. We also made a third set of constructs that added both the chick/quail RUNX2 SNPs and SMAD binding element to the duck *Mmp13* promoter and vice versa. We cloned each of these −184/181 fragments into a luciferase expression vector and transfected them into chick and duck cells, which were then treated with rTGFβ1.

**Figure 7.**
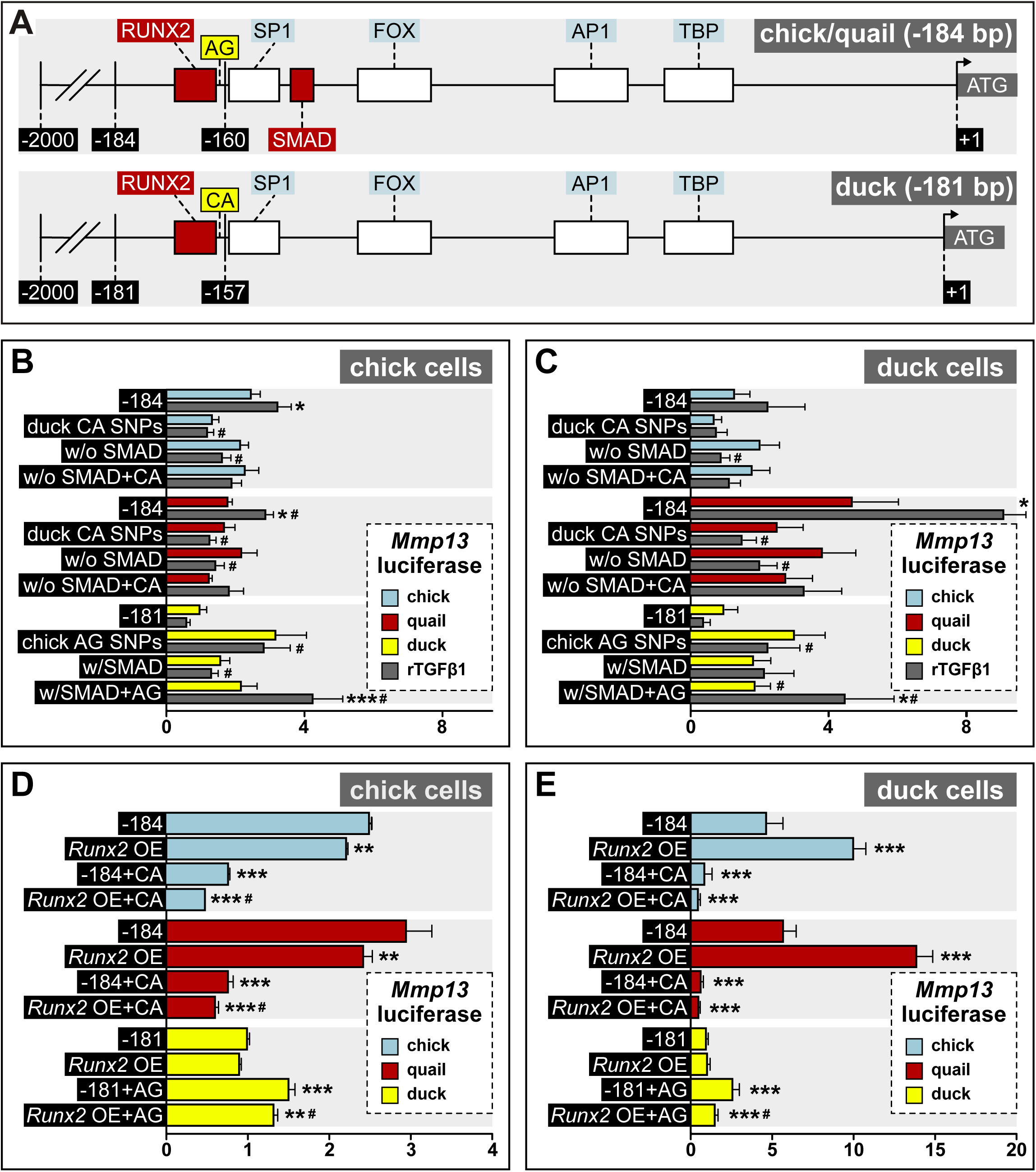
Response of the *Mmp13* promoter to rTGFβ1 in relation to species-specific SNPs and RUNX2 and SMAD binding elements. **(A)** Schematic of the proximal region of the chick/quail (−184 bp) and duck (−181 bp) promoters. Chick/quail contain 2 SNPs (“AG”) adjacent to a RUNX2 binding element that differ from duck (“CA”). **(B)** Chick and **(C)** duck cells transfected with the chick (blue) or quail (red) *Mmp13* promoter containing either a 184 bp control fragment including the chick/quail “AG” SNPs plus one RUNX2 and one SMAD binding element; a 184 bp fragment including the duck “CA” SNPs plus one RUNX2 and one SMAD binding element; a 184 bp fragment including the chick/quail “AG” SNPs plus one RUNX2 but without one SMAD binding element; or a 184 bp fragment including the duck “CA” SNPs plus one RUNX2 but without one SMAD binding element. Chick and duck cells were also transfected with the duck (yellow) *Mmp13* promoter containing either a 181 bp control fragment including the duck “CA” SNPs plus one RUNX2 but no SMAD binding element; a 181 bp fragment including the chick/quail “AG” SNPs plus one RUNX2 but no SMAD binding element; a 181 bp fragment including the duck “CA” SNPs plus one RUNX2 and one SMAD binding element; or a 181 bp fragment including the chick/quail “AG” SNPs plus one RUNX2 and one SMAD binding element. Cells were treated with rTGFβ1 (dark gray) and assayed for luciferase activity. Switching the chick/quail SNPs to “CA” SNPs abolishes rTGFβ1 induction. Switching the duck SNPs to “AG” increases basal activity but does not add a rTGFβ1 response. Removing the chick/quail SMAD binding element abolishes rTGFβ1 induction. Switching both the RUNX2 SNPs and SMAD binding element in chick/quail with the duck sequence abolishes rTGFβ1 induction. Switching both the AG SNPs and the SMAD binding element into the duck promoter adds a response to rTGFβ1 (n = 24). * denotes significance from untreated control within each group p < 0.05, ** denotes significance from empty vector control within each group p < 0.01, *** denotes significance from empty vector control within each group p < 0.001, # denotes significance between quail and duck, † denotes significance between chick and quail, ‡ denotes significance between chick and duck. **(D)** Chick and **(E)** duck cells transfected with the chick or quail *Mmp13* promoter containing either a 184 bp control fragment including the chick/quail “AG” SNPs; a 184 bp fragment including the chick/quail “AG” SNPs plus a *Runx2* over-expression (OE) construct; a 184 bp fragment including the duck “CA” SNPs; or a 184 bp fragment including the duck “CA” SNPs plus a *Runx2* OE construct. Chick and duck cells were also transfected with the duck *Mmp13* promoter containing either a 181 bp control fragment including the duck “CA” SNPs; a 181 bp fragment including the duck “CA” SNPs plus a *Runx2* OE construct; a 181 bp fragment including the chick/quail “AG” SNPs; or a 181 bp fragment including the chick/quail “AG” SNPs plus a *Runx2* OE construct. In chick cells, *Runx2* OE decreased *Mmp13* promoter activity for both chick and quail promoters, while no change was observed in the duck promoter. Putting the duck SNPs in chick/quail leads to a decrease in *Mmp13* promoter activity even with *Runx2* OE. Putting the chick/quail SNPs in duck leads to a decrease in promoter activity with *Runx2* OE. In duck cells, *Runx2* overexpression induced *Mmp13* promoter activity in chick and quail, while no change was observed in the duck promoter. Switching the SNPs in chick/quail to duck abolished the *Runx2* induction. However, switching the SNPs in duck to chick/quail repressed promoter activity with Runx2 overexpression (n = 24). * denotes significance from the endogenous promoter within each species p < 0.05, ** denotes significance the endogenous promoter within each species p < 0.01, *** denotes significance from the endogenous promoter within each species p < 0.001, # denotes significance between control SNP switch versus *Runx2* overexpression with the SNP switch within each species.

We find that putting the duck “CA” SNPs into the chick and quail *Mmp13* promoters lowers activity and abolishes induction by rTGFβ1 (Figure 7B-C). Similarly, removing the SMAD binding element reduces chick and quail promoter activity and diminishes the response to rTGFβ1. Moreover, chick and quail promoters containing both the duck “CA” SNPs and SMAD binding element deletion show an equivalent diminished response to rTGFβ1 compared to the −184 fragment treated with rTGFβ1. In contrast, when we put the chick/quail “AG” SNPs into the duck promoter, we observe an increase in endogenous activity, although the SNP switch alone is not sufficient to recover induction by rTGFβ1. We observe the same effect with increased endogenous activity when we put the SMAD binding element into the duck promoter, but no induction with rTGFβ1. However, when we put both the chick/quail “AG” SNPs and the SMAD binding element into the duck promoter, we increase endogenous activity and trigger sensitivity to rTGFβ1, with a 2-fold induction in chick cells (p < 0.002) and 2.4-fold induction (p < 0.05) in duck cells compared to the untreated promoter (Figure 7B-C).

To test if the SNPs adjacent to the RUNX2 binding element play a role in the species-specific regulation of the *Mmp13* promoter by RUNX2, we transfected chick and duck cells with the −181/184bp *Mmp13* promoter constructs and with constructs in which we switched the chick/quail “AG” and duck “CA” SNPs. These same chick and duck cells were also co-transfected with either an empty vector or a dox-inducible *Runx2* over-expression construct (Chu et al., 2020). Our results show that in chick cells, *Runx2* over-expression significantly decreases the activity of the chick and quail *Mmp13* promoter, whereas *Runx2* over-expression has little effect on the duck promoter (Figure 7D). In chick cells, replacing the chick/quail SNPs with the duck SNPs decreases activity of the chick (3.2-fold, p < 0.0001) and quail (3.8-fold, p < 0.0001) promoters, and *Runx2* overexpression reduces promoter activity in quail (1.2-fold, p < 0.001) but not chick. Replacing the duck SNPs with chick/quail SNPs induces activity of the promoter by 1.5-fold (p < 0.0001), and *Runx2* overexpression does not change *Mmp13* promoter activity. In duck cells, *Runx2* overexpression increases *Mmp13* promoter activity for chick (1.9-fold, p < 0.0001) and quail (1.9-fold, p < 0.0001), while causing no change in activity for the duck promoter (Figure 7E). Replacing the chick/quail SNPs with the duck SNPs decreases promoter activity for chick (7.2-fold, p < 0.0001) and quail (8.9-fold, p < 0.0001), and the induction observed from *Runx2* over-expression in the endogenous promoter is abolished when the SNPs are switched. Replacing the duck SNPs with the chick/quail SNPs increases activity of the promoter by 2.6-fold (p < 0.0001), and *Runx2* over-expression decreases promoter activity by 1.7-fold (p < 0.003).

## DISCUSSION

### MMP13 is co-expressed with TRAP and shows species-specific regulation

Previously, we demonstrated that *Mmp13* and bone resorption are regulated by NCM and are important for controlling the species-specific length of the jaw skeleton (Ealba et al., 2015). Briefly, we observed that quail have higher levels of bone resorption markers than do duck during jaw development, including TRAP, and this difference increases over time. Moreover, transplanting NCM from quail to duck dramatically elevates expression of *Mmp13* and TRAP and generates chimeras with shorter quail-like jaws. In these chimeras, the *Mmp13*-expressing cells arise from the quail-donor NCM. In this prior work, we quantified the ratio of TRAP to bone (*i.e.,* osteoid) volume in duck, quail, and chimeric quck. We found significant differences between duck and quail in the amount of bone resorption, and that the donor side of quck is more quail-like whereas the host side is more duck-like. Moreover, we showed that blocking resorption using a bisphosphonate or an MMP13 inhibitor significantly lengthens the jaw whereas activating bone resorption can shorten the jaw (Ealba et al., 2015).

A goal of the present study was to identify regulatory mechanisms that lead to the significantly higher levels of *Mmp13* expression and bone resorption observed in quail relative to duck during development of the lower jaw. Our IHC analyses reveal that wherever bone resorption is elevated in the jaw, so is MMP13, and that there are clear differences between quail and duck in both the overall levels and spatial domains of expression. MMP13 is a type 1 collagenase involved in bone resorption and is produced by osteoblasts, osteocytes, as well as by mature chondrocytes (Johansson et al., 1997; Sasano et al., 2002; Hatori et al., 2004; Stickens et al., 2004; Holmbeck et al., 2005; Behonick et al., 2007; Tang et al., 2012; Zhang et al., 2012; Yamamoto et al., 2016). In the jaw skeleton, all of these *Mmp13*-expressing cell types are derived entirely from NCM (Le Lièvre, 1978; Noden, 1978; Helms and Schneider, 2003). Although *Mmp13* is expressed by hypertrophic chondrocytes when cartilage is replaced by bone during endochondral ossification (Colnot and Helms, 2001), during development of the lower jaw in birds, Meckel’s cartilage persists and there is no endochondral ossification except for that limited entirely to the most proximal region within the articular cartilage beginning at HH39 (Starck, 1989; Eames et al., 2004; Mitgutsch et al., 2011; Svandova et al., 2020). All other bones in the lower jaw including the angular and the dentary, which were a focus in our study, form through intramembranous ossification and undergo perilacunar remodeling (Helms and Schneider, 2003).

Overall, we observe higher levels of TRAP staining in bones of the quail lower jaw compared to those in stage-matched duck. Specifically, when we examined the quail angular bone, we find elevated MMP13 protein levels coincident with greater amounts of TRAP staining, whereas in the angular bone of duck we observe very low levels of TRAP and MMP13 despite substantial amounts of osteoid. Within the dentary bone of both quail and duck, however, MMP13 and TRAP are co-expressed, although levels for quail appear much more elevated than those in duck. This finding indicates there are regulatory mechanisms that can be deployed spatially and control both the specific-specific levels and bone-specific domains of MMP13 and TRAP expression, which ultimately may create zones of remodeling that regulate the size and shape of the jaw skeleton. Such a result is consistent with other work proposing that differential fields of resorption underlie changes in the size and shape of the developing human jaw skeleton (Enlow et al., 1975; Moore, 1981; Radlanski and Klarkowski, 2001; Radlanski et al., 2004).

### TGFβ signaling mediates species-specific bone resorption in the jaw skeleton

TGFβ signaling is involved in a diverse array of cellular processes, such as promoting cell proliferation, cell survival, migration, and production of extracellular matrix (ECM) (Stouffer and Owens, 1994; Moses and Serra, 1996; Massague and Wotton, 2000; Vinals and Pouyssegur, 2001; Verrecchia and Mauviel, 2002; Derynck et al., 2008; Fang et al., 2012). We focused on TGFβ signaling because this pathway is known to be expressed in craniofacial tissues during avian development (Yamagishi et al., 1999; Cooley et al., 2014; Woronowicz et al., 2018) and affect jaw length based on our own work on TGFβ targets such as *Mmp13* and *Runx2* (Hall et al., 2014; Ealba et al., 2015) and from studies on mouse and human mutations (Ito et al., 2003; Dudas et al., 2006). For example, mutations in *Tgfβ2* and *Tgfβr1* cause microretrognathia in Loeys-Dietz Syndrome (Loeys et al., 2005; Zhao et al., 2008) whereas elevated TGFβ signaling in Marfan syndrome causes excessive upper jaw growth (Westling et al., 1998; Neptune et al., 2003). Jaw length defects are also associated with mutations in *Tgfβr2* and *Smad2* (Nomura and Li, 1998; Oka et al., 2007; Oka et al., 2008), and dysregulation in noncanonical NCM-mediated TGFβ signaling results in hypoplastic facial features (Yumoto et al., 2013). But exactly how such changes to members and targets of the TGFβ pathway can ultimately modulate jaw length remain unclear.

TGFβ signaling plays a crucial role in the local osteocyte-mediated perilacunar remodeling of bone (Dole et al., 2017). Thus, we tested if species-specific levels of bone resorption and *Mmp13* expression observed in quail versus duck correlate with differential regulation of the TGFβ pathway. We quantified expression of TGFβ pathway members and targets in the developing jaw primordia of quail, chick, and duck, and find higher levels of *Tgfβ1, Tgfβ3,* and *Tgfβr1* in quail and chick versus duck at HH37, which is the stage when bone resorption can first be detected via TRAP staining in the lower jaw (Ealba et al., 2015). To confirm activation of the TGFβ pathway, we assayed for pSMAD2 and pSMAD3 by western blot and observe higher levels in quail and chick versus duck at HH37, indicating that quail have elevated TGFβ signaling. Quail also show higher expression of target genes including *Runx2*, *Mmp13*, *Pai1*, and *Mmp2*. At a slightly later stage of quail development (*i.e.,* HH40) when bone mineralization and resorption are elevated (Hall et al., 2014; Ealba et al., 2015), we observe that *Tgfβ3, Smad2,* and *Mmp13,* as well as pSMAD3 decrease in expression. This implies that once the TGFβ pathway becomes activated and initiates bone resorption, there is negative feedback to ensure dampened signaling at later stages. This observation is supported by multiple studies finding both dose and time to be crucial parameters in mediating and determining the effects of TGFβ (Alliston et al., 2001; Takeuchi et al., 2010; Chen et al., 2012a; Abou-Ezzi et al., 2019). Notably, we do not observe differences in the expression of all TGFβ pathway members among chick, quail, and duck, including *Tgfβ2, Tgfβr2*, *Tgfβr3,* or *Smad3*, suggesting that species-specific regulation of the pathway may be facilitated by certain genes and not others.

To test for a mechanistic connection between TGFβ signaling and bone resorption, we treated lower jaws from quail with TGFβR1, SMAD3, or MMP13 inhibitors and then assayed for changes in bone resorption via TRAP staining. We find that inhibiting the TGFβ pathway at the level of a receptor, an intracellular mediator, or a target gene leads to a decrease in bone resorption. This supports our published work showing a direct link between the species-specific amount of bone resorption and jaw length (Ealba et al., 2015), and reveals that TGFβ signaling is an indispensable determinant. Our findings are also consistent with a previous study showing that TGFβ signaling plays an important role in the local osteocyte-mediated perilacunar remodeling of bone (Dole et al., 2017). Thus, the elevated levels of bone resorption observed in the jaw skeleton of quail relative to duck depend upon TGFβ signaling.

Additionally, we find that chick show a similar stage-specific expression profile as quail, with increases in *Tgfβ1*, *Tgfβ3, Smad2, Runx2, Pai1, Mmp2, Mmp9,* and *Mmp13* during the period associated with bone resorption. However, chick tends to have absolute levels of gene expression intermediate to quail and duck, which may reflect the fact that the chick jaw skeleton falls somewhere in-between the size of quail and duck (Schneider and Helms, 2003; Eames and Schneider, 2008; Mitgutsch et al., 2011). For *Smad2* and *Smad3,* chick had higher levels of expression compared to quail and duck, but this was not correlated with higher pSMAD3 activation, perhaps due to post-transcriptional regulation of *Smad2* and *Smad3*. Taken together, our comparative analysis of the TGFβ pathway in chick, quail, and duck demonstrates that certain ligands, receptors, intracellular mediators, and downstream effectors are upregulated and activated in quail and chick lower jaws when bone resorption is initiated, and this could potentially be one mechanism whereby quail and chick achieve higher levels of bone resorption and generate a relatively shorter jaw length than duck during development.

### Sensitivity to TGFβ and differential activation of target genes are species-specific

Our study reveals that the developing lower jaw of duck shows significantly less sensitivity to TGFβ signaling than the developing lower jaws of chick and quail, which respond with greater pSMAD3 activation and induction of *Runx2* and *Mmp13* when treated with rTGFβ1. We observe equivalent differences in the species-specific response of duck fibroblasts when compared to chick fibroblasts, indicating that sensitivity to TGFβ signaling is cell autonomous, and suggesting that phosphorylation of SMAD3 and the induction of *Runx2* and *Mmp13* in quail versus duck does not depend upon the context of development. This does not mean that duck cells are unable to respond to rTGFβ1, since we observe faster and greater induction of other TGFβ targets including *Pai1* and *Mmp2* when compared to chick cells (although we do not observe such a response in duck lower jaws). We also find that rTGFβ1 has no measurable effect on the expression of bone differentiation markers including *Col1a1* in any species, or on *Ocn* in chick and quail. Treatment with rTGFβ1 appears to repress *Ocn* in duck. Previous work has demonstrated that the induction or repression of these genes depends upon the stage of differentiation as well as other factors (Garcia-Trevijano et al., 1999; Palcy et al., 2000; Gurlek and Kumar, 2001; Subramaniam et al., 2001; Iwata et al., 2010; Chen et al., 2012b; Pan et al., 2013). In our experiments, lower jaws were cultured and treated just prior to the beginning of mineralization, which may have not been the most appropriate timepoint for assaying changes to expression of such late-stage osteogenic markers. Moreover, these lower jaws could require a longer time in culture to differentiate further, and thus 24 hours of treatment might not have been sufficient to observe any effect.

Nonetheless, the finding that some pathway targets are responsive to rTGFβ1 and others are not, provides a valuable internal control and indicates that additional regulatory mechanisms are at work that underlie the differential sensitivity to TGFβ signaling and enable the species-specific activation of *Runx2* and *Mmp13* in quail versus duck. Potential mechanisms could involve feedback upon the pathway itself and/or genetic/epigenetic changes to the regulatory landscape especially at the level of target genes. As a proof-of-concept, we explored such possibilities by examining the effects of treatments on TGFβ receptor expression and by focusing on species-specific variation in SMAD and RUNX2 binding elements in the *Mmp13* promoter. In terms of the effects of TGFβ signaling on receptor expression we observed no changes in *Tgfβr1*, *Tgfβr2*, or *Tgfβr3* expression in lower jaws treated with rTGFβ1. This suggests that any positive or negative feedback on pathway activation or on levels of receptor expression may occur independent of ligand availability, which differs from what has been observed in another system (Duan and Derynck, 2019). That being said, our inhibitor experiments further demonstrate that the ability of rTGFβ1 to induce *Runx2* and *Mmp13* occurs directly via activation of TGFβR1 and phosphorylation of SMAD3, which is something that has been observed previously (Selvamurugan et al., 2004a; Chen et al., 2012b; Di Chen, 2019). Although inhibition of TGFβR1 can lead to upregulation of *Runx2* and *Mmp13* via a non-canonical p38 MAPK pathway (Chen et al., 2012a), which is something we did not analyze in the current study.

In this context, one scenario to account for differences in sensitivity to rTGFβ1 between species could be the differential regulation of *Tgfβr1*, since we observe significantly higher levels of expression in quail during three of the four developmental stages analyzed. TGFβR1 is a transmembrane serine/threonine kinase that heterodimerizes with TGFβR2 and is a critical component of TGFβ signal transduction (Cheifetz et al., 1990; Laiho et al., 1990; Franzen et al., 1993; Tomoda et al., 1994; Feng and Derynck, 1997; Vellucci and Reiss, 1997; Chen et al., 2006; Derynck et al., 2008). TGFβR1 is activated by transphosphorylation once TGFβ ligand binds to TGFβR2, which in turn activates target genes (Wrana et al., 1994; Massague and Wotton, 2000). Since we observed constant and comparable levels of *TGFβR2* expression in chick, quail, and duck, and TGFβR1 must heterodimerize with TGFβR2 in order for signal transduction to occur, then a rate determining step that could serve as a mechanism for modulating pathway activation could be the species-specific regulation of *Tgfβr1*. Determining how *Tgfβr1* is differentially regulated between quail and duck remains a subject of interest for future research. SNPs in *Tgfβr1* and in its promoter are linked to decreased *Tgfβr1* expression as well as craniofacial, skeletal, and other disorders including jaw length defects (Wu et al., 2002; Chen et al., 2004; Loeys et al., 2005; Pasche et al., 2005; Zhao et al., 2008; Pasche et al., 2010; Sun et al., 2011; Knobloch et al., 2019). Receptor localization may also play a vital role since previous studies have shown that cells can modulate the translocation of TGFβ receptors from the cytosol to their integration at the plasma membrane and allow ligands to bind and initiate signal transduction (Zhang et al., 1999; Rys et al., 2015; Duan and Derynck, 2019). TGFβ receptors can become internalized through clathrin-dependent endocytosis or lipid rafts facilitated by caveolin-1 (Anders et al., 1998; Di Guglielmo et al., 2003; Luga et al., 2009). Once internalized, they can further induce signal transduction, be recycled, or be targeted for degradation (Mitchell et al., 2004; Penheiter et al., 2010). Additional work could involve comparing dimerization and translocation of TGFβ receptors at the protein level in quail versus duck.

### Promoter evolution underlies the species-specific expression of Mmp13

To identify molecular mechanisms underlying the species-specific expression of *Mmp13,* we focused on the structure and function of the *Mmp13* promoter. The *Mmp13* promoter contains SMAD binding elements and other domains that regulate its expression (Pendas et al., 1997; Wang et al., 2004; Chen et al., 2012a; Meyer et al., 2016; Takahashi et al., 2017; Young et al., 2019). We observed that a 2 kb fragment of the quail and chick *Mmp13* promoter can be induced by rTGFβ1, but not the 2 kb fragment of the duck *Mmp13* promoter. Additionally, *Mmp13* is a target of RUNX2, and RUNX2 can have both an activating and repressive effect on *Mmp13* expression depending on the context and the duration of treatment (Chen et al., 2012a). We find that *Runx2* is both more highly expressed in the lower jaws of quail and much more activated in response to treatments with rTGFβ1 than in those of duck. Therefore, we also performed overexpression experiments to test the extent to which *Runx2* can activate the duck *Mmp13* promoter. While the −2 kb promoter of quail shows a strong induction following *Runx2* over-expression, the duck promoter shows no response and trends towards repression. This suggests the presence and/or absence of regulatory elements within the *Mmp13* promoter of chick, quail, and duck that facilitate the species-specific response to TGFβ signaling via the differential binding of SMAD and/or RUNX2.

Our comparative analysis of the *Mmp13* promoter reveals that along a −2 kb fragment upstream from the transcriptional start site there are five SMAD binding elements in chick, four in quail, and three in duck; and three RUNX2 binding elements in chick, quail, and duck. Not only are these binding elements found at different sites across the *Mmp13* promoter in each species, but also one of these SMAD binding elements is present in the most proximal portion of the chick and quail proximal promoter but absent in duck. This particular SMAD binding element has been shown to be regulated by SMAD3 in the human and mouse *Gsc* promoter (Martin-Malpartida et al., 2017). To test if the species-specific response to TGFβ signaling and expression of *Mmp13* is mediated by these SMAD and RUNX2 binding elements, we generated different sets of reporter constructs with or without these binding sites in the *Mmp13* promoter for each species. We transfected chick and duck cells with each of these constructs and treated them with rTGFβ1.

Overall, we find that removing the most proximal SMAD binding element reduces the response in quail and chick but adding this binding element to the duck promoter increases overall activity. We also identified SNPs directly adjacent to a RUNX2 binding site that distinguish quail and chick (*i.e.*, “AG”) from duck (*i.e.,* “CA”). We deduce that these SNPs may play an essential role in RUNX2 binding because switching them along with the SMAD binding element between species abolishes the response to rTGFβ1 with quail and chick promoters but adds a response to duck. Altering one of these individual elements alone is not sufficient to induce a rTGFβ1 response, suggesting that cooperativity at the SMAD binding element plus the RUNX2 binding element when “AG” SNPs are present is necessary to achieve induction. Other studies have shown that such cooperative binding and modifications to various sites can affect *Mmp13* promoter activation, including those for methylation of *Hif1α*, and binding of YP-1, LEF1, osterix, vitamin D receptor, FOS, JUN, and PTH (Samuel et al., 2007; Yun and Im, 2007; Hashimoto et al., 2013; Meyer et al., 2016). Our findings provide additional evidence that the “AG” SNPs and SMAD binding element work cooperatively and are required for TGFβ activation of the *Mmp13* promoter, and that species-specific expression levels are mediated by the structure of the promoter. In this regard, evolutionary changes within the promoter of chick, quail, and duck appear to be a central mechanism that controls the species-specific expression of *Mmp13*.

## CONCLUSION

In “*Problems of Relative Growth*” Huxley (1932) proposed genetic mechanisms for generating phenotypic diversity that included mutations affecting what he called time and rate genes. Huxley’s critical insight was that these types of mutations could regulate where and when a given gene turns on or off and titrate its expression, which ultimately would then alter growth parameters and anatomy at many levels in coordinated ways (Schneider, 2018a). In this regard, regulatory changes have been viewed as a primary mechanism for evolutionary diversification instead of modifications to coding sequences of genes (Britten and Davidson, 1969; King and Wilson, 1975; Carroll, 2005; Wray, 2007; Romero et al., 2012). Given that roughly 98% of DNA in the human genome is non-coding (Clamp et al., 2007), deciphering the morphogenetic consequences of mutations in regulatory domains is necessary to illuminate fundamental mechanisms of development, disease, and evolution (Ahituv, 2012). While many studies have focused on *cis*-regulatory enhancers, which are often located tens to hundreds of kb from transcriptional start sites (Heintzman et al., 2007; Shen et al., 2012; Prescott et al., 2015; Schaffner, 2015; Long et al., 2016; Rebeiz and Tsiantis, 2017; Williams et al., 2018; Kim et al., 2019), there are some examples of how evolution within the more proximal promoter itself can drive phenotypic change. The globin genes are a well-studied case (Pace and Makala, 2012), although these also appear to rely on distal enhancers for their regulation (Hay et al., 2016). When broadly comparing non-coding regulatory regions across species, enhancers appear to undergo rapid evolutionary turnover whereas promoters tend to remain partially or fully conserved (Villar et al., 2015; Berthelot et al., 2018). MMP family members are no exception and many share structural similarity and numerous conserved binding elements in their proximal promoters (Benbow and Brinckerhoff, 1997; Samuel et al., 2007; Yan and Boyd, 2007; Fanjul-Fernandez et al., 2010; Hashimoto et al., 2013). For this reason, we think our finding that species-specific changes in the *Mmp13* promoter, which alter its sensitivity to TGFβ signaling and its regulation by RUNX2, offers a novel insight on a potential developmental mechanism for generating evolutionary variation in jaw length. Similarly, polymorphisms in the *Mmp13* promoter that affect binding of transcriptional activators or repressors can generate abnormal variation associated with human disease (Pendas et al., 1997; Ye, 2000; Marchenko et al., 2002; Yoon et al., 2002; Achari et al., 2008).

While our results indicate that promoter evolution may play an important role in the species-specific expression of *Mmp13* and in this way could influence the establishment of jaw length in quail versus duck via bone resorption, the potential contributions from additional levels of *Mmp13* regulation, multiple *Mmps*, as well as different members and targets of TGFβ and other signaling pathways remain vast (Mina, 2001b; Mina, 2001a; Depew et al., 2002; Mina et al., 2002; Chakraborti et al., 2003; Oka et al., 2007; Havens et al., 2008; Balic et al., 2009; Fanjul-Fernandez et al., 2010; Fish et al., 2011; Young et al., 2019). In prior studies we have found that NCM-mediated programs underlying jaw development in quail and duck also vary due to species-specific differences in the cell biological properties of NCM and in the physical and signaling interactions between NCM and adjacent tissues (Schneider and Helms, 2003; Schneider, 2007; Eames and Schneider, 2008; Merrill et al., 2008; Tokita and Schneider, 2009; Solem et al., 2011; Fish and Schneider, 2014b; Fish et al., 2014; Hall et al., 2014; Ealba et al., 2015; Schneider, 2015; Woronowicz et al., 2018; Woronowicz and Schneider, 2019).

Likewise, we predict that species-specific variation in cis-regulatory domains at more distal enhancers, in the complement of available transcriptional cofactors, in epigenetic mechanisms of transcriptional and post-transcriptional control such as DNA methylation and non-coding RNA, in the post-translational modification of proteins, and in the gradients and thresholds of secreted molecules will similarly contribute in some meaningful manner (Schneider, 2018b; Schneider, 2018a). By focusing on species-specific regulation of members and targets of the TGFβ pathway in the avian jaw, we hope our work has helped pinpoint precisely when and where one mode of change in a developmental program can alter the course of morphological evolution.

## ACKNOWLEDGEMENTS

We thank T. Alliston and R. Marcucio for helpful discussions and comments on the manuscript; J. Aggleton, Z. Vavrušová, A. Nguyen, A. Lucena, G. Krish, P. Asfour, and T. Huang for technical assistance. We thank T. Dam at AA Lab Eggs. The pmScarlet-i_C1 was a gift from Dorus Gadella (Addgene, #85044). The AAVS1 Puro Tet3G 3xFLAG Twin Strep was a gift from Yannick Doyon (Addgene, # 92099). The pKanCMV-mClover3-mRuby3 was a gift from Michael Lin (Addgene, #74252). The pCAG-Cre-IRES2-GFP was a gift from Anjen Chenn (Addgene, #26646). The pCMV-hyPBase was provided by the Wellcome Trust Sanger Institute. The mNeonGreen was provided by Allele Biotechnology & Pharmaceuticals. This work was supported in part by the UCSF Biological Imaging Development Core (BIDC); the UCSF Core Center for Musculoskeletal Biology and Medicine (CCMBM) through NIAMS P30 AR066262; NIDCR F31 DE027283 to S.S.S.; and NIDCR R01 DE016402 and R01 DE025668, and NIH Office of the Director S10 OD021664 to R.A.S.

---

**Supplemental Figure 1. Sequence logos for RUNX2 and SMADs.** Graphical representation of the binding profiles of the position weight matrices taken from the JASPAR 2020 database for **(A)** RUNX2, **(B)** SMAD2/3 heterodimer, **(C)** SMAD3, and **(D)** SMAD4. The relative sizes of the letters indicate their frequency in the sequences whereas the height of the letters shows the information content of the position.

**Supplemental Figure 2. Relative mRNA of TGFβ pathway members and targets in developing lower jaws of chick, quail, and duck. (A)** RT-qPCR analyses reveal *Tgfβ2* mRNA expression does not change in chick (blue circles; n = 8) and quail (red squares; n = 8) over developmental time, however in duck (yellow triangles; n = 10) there is a 2.2 and 3.3-fold decrease at HH37 and HH40; *Tgfβ2* expression is significantly higher in duck than chick or quail at HH31. **(B)** *Acvrl1* expression does not change over time in chick, however quail show a 3.9-fold increase at HH37. There is a 1.7-fold reduction in duck at HH40. Duck has significantly higher *Acvrl1* expression at HH34 compared to chick and quail. However, at HH37 and HH40 quail has significantly more *Acvrl1* expression compared to chick or duck. **(C)** *Tgfβr2* mRNA expression does not significantly change over time in chick or duck, but in quail there is a 1.8-fold induction at HH37. No significant differences are observed between the species. **(D)** *Tgfβr3* mRNA expression does not change over time in chick, quail, or duck. No significant differences are observed between species. **(E)** *Smad3* mRNA expression in chick does not change until HH40 with a 2.7-fold reduction. In quail and duck, *Smad3* expression decreases at HH37 and continues to decrease for duck at HH40. Chick *Smad3* expression is significantly higher than quail at HH34 and is significantly higher at HH37 compared to quail and duck. **(F)** *Pai1* mRNA expression does not change over time in chick, but quail increases 2.6-fold at HH37 and HH40, while duck decreases 4.7-fold at the same time points. Duck *Pai1* expression is significantly higher at HH31 and HH34 compared to chick and quail, but at HH37 and HH40 quail it is significantly higher compared to chick and duck. **(G)** *Mmp2* mRNA expression increases 2-fold in chick and 4.7-fold in quail at HH37, whereas duck decreases 10-fold at HH40. Chick is significantly higher in *Mmp2* expression compared to duck at HH34 and HH37. Quail has substantially higher *Mmp2* expression compared to duck at HH37 and HH40. **(H)** *Mmp9* mRNA expression increases 64-fold in chick, 30-fold in quail, and 43-fold in duck at HH37 and maintains similar levels of induction at HH40. Quail has significantly more *Mmp9* expression compared to chick and duck at HH37 and HH40. p < 0.05 and * denotes significance from HH31 within each group, # denotes significance between quail and duck at same stage, † denotes significance between chick and quail, ‡ denotes significance between chick and duck.

**Supplemental Figure 3. Representative western blot images.** Western blot images for **(A)** pSMAD3 and **(B)** MMP13 in chick (n = 9), quail (n = 12), and duck (n = 12) jaws from stages HH31-HH40 with β-Actin as a loading control. Dashed lines represent images taken from 2 separate fluorescent channels on the same gel, 680RD for β-Actin and 800CW for pSMAD3 or MMP13. **(C)** pSMAD3 and **(D)** MMP13 in chick (DF-1) and duck (CCL-141) cells treated with 5 ng/ml rTGFβ1 for 2 hours with β-Actin as a loading control (n = 8). **(E)** RUNX2 and **(F)** MMP13 levels in chick cells treated for 24 hours with rTGFβ1, TGFβR1 inhibitor, a combination of both rTGFβ1 and TGFβR1 inhibitor, SMAD3 inhibitor, and a combination of both rTGFβ1 and SMAD3 inhibitor (n = 8). **(G)** pSMAD3 and **(H)** MMP13 levels in chick, quail, and duck lower jaws treated in culture with 10, 25, and 50 ng/ml of rTGFβ1 for 6 hours or 24 hours respectively (n = 12).

**Supplemental Figure 4. Relative mRNA for markers of bone remodeling in the developing lower jaws of chick, quail, and duck. (A)** RT-qPCR analyses reveals *Mmp14* mRNA expression does not change in chick (n = 8) and quail (n = 8), or duck (n = 10). **(B)** *Cathepsin K (Ctsk)* increases in chick 1.5-fold, and 4-fold in quail at HH37. There is no change in expression in duck. There is a significantly higher expression in quail compared to duck at HH37. **(C)** *Sost* increases 3.6-fold in chick and quail at HH37. Duck *Sost* expression does not change over developmental time. Chick and quail have significantly higher *Sost* expression at HH37 and HH40 compared to duck at comparable stages. p < 0.05 and * denotes significance from HH31 within each group, # denotes significance between quail and duck at same stage, † denotes significance between chick and quail, ‡ denotes significance between chick and duck.

**Supplemental Figure 5. Sensitivity of chick and duck cells, as well as chick, quail, and duck lower jaws to rTGFβ1 on TGFβ target genes and *Tgfβr* expression. A)** *Pai1* expression in chick (blue) and duck (yellow) cells treated for 1 to 24 hours with rTGFβ1 (dark gray). rTGFβ1 induces chick *Pai1* mRNA expression by 3.7-fold at 24 hours, while duck *Pai1* increases 2-fold at 1 hour and 2.4-fold at 24 hours (n = 8). **(B)** *Mmp2* expression increases in chick cells 1.9 and 2-fold with rTGFβ1 treatment at 6 and 24 hours respectively, while in duck cells *Mmp2* is induced 3.5 and 7.5-fold at 1 and 24 hours. In rTGFβ1 treated cells, duck shows significantly higher *Mmp2* expression at 1 and 24 hours compared to chick (n = 8). **(C)** Chick (blue), quail (red), and duck (yellow) lower jaws treated in culture with 25 ng/ml (dark gray) of rTGFβ1 for 24 hours. *Tgfβr1* expression does not significantly change with rTGFβ1in any species (n = 10). (D) *Tgfβr2* expression does not significantly change with rTGFβ1in any species (n = 10). **(E)** Chick and duck *Pai1* expression does not change with rTGFβ1 treatment, whereas there is a 1.5-fold induction in quail. Quail and chick show significantly higher *Pai1* expression compared to duck (n = 10). (F) *Mmp2* expression increases 1.3 and 1.4-fold with rTGFβ1 treatment in chick and quail lower jaws, while duck does not change. Quail and chick has significantly higher *Mmp2* expression compared to duck (n = 10). **(G)** Collagen 1A1 (*Col1a1)* mRNA expression does not change significantly with rTGFβ1 treatment in any species (n = 10). (H) *Osteocalcin (Ocn)* expression does not change significantly with rTGFβ1 treatment in any species (n = 10). * denotes significance from HH31 within each group p < 0.05, ** denotes significance from HH31 within each group p < 0.01, # denotes significance between quail and duck at same stage, † denotes significance between chick and quail, ‡ denotes significance between chick and duck.

