## Supplemental Figures for "Species-specific sensitivity to TGFβ signaling and changes to the Mmp13 promoter underlie avian jaw development and evolution"

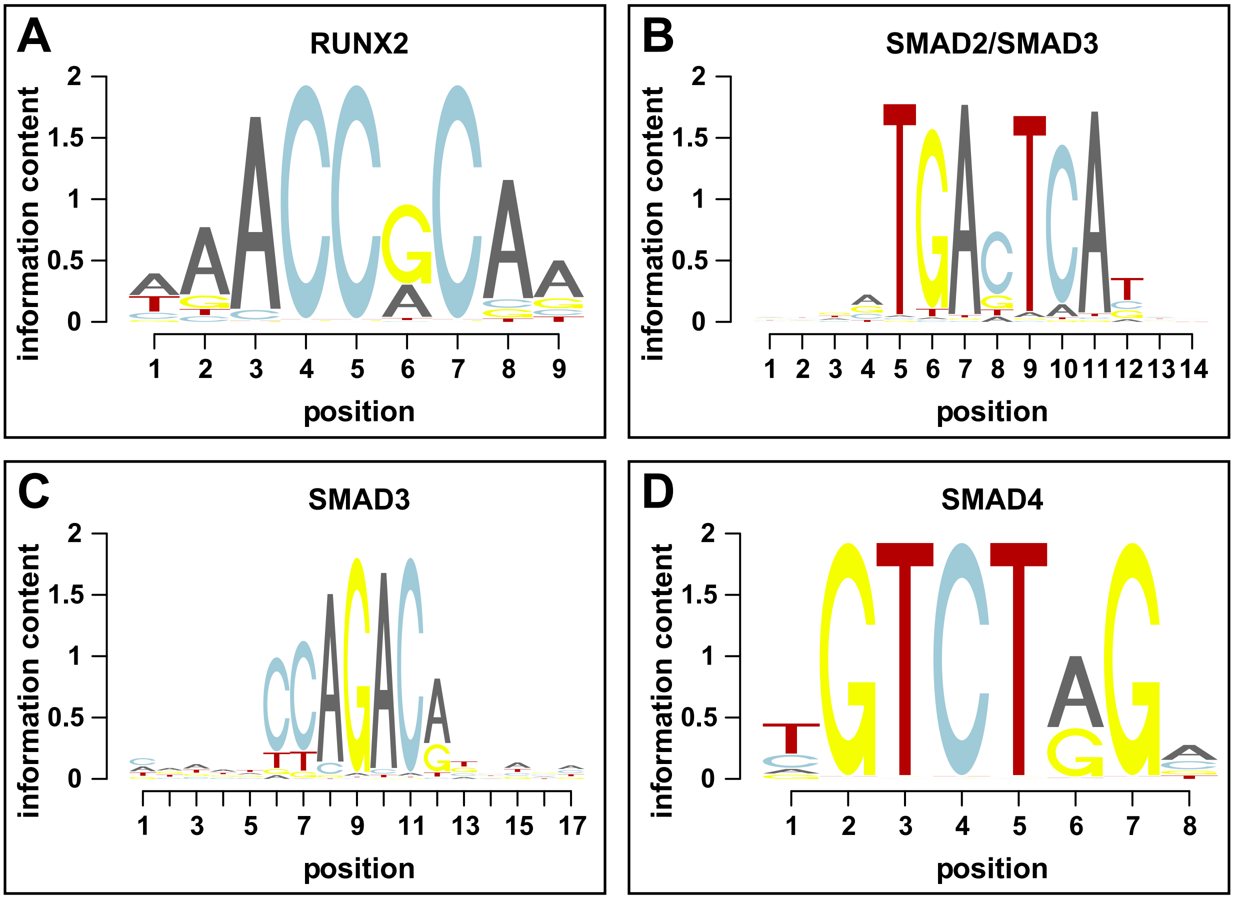


**SUPPLEMENTAL FIGURE 1**

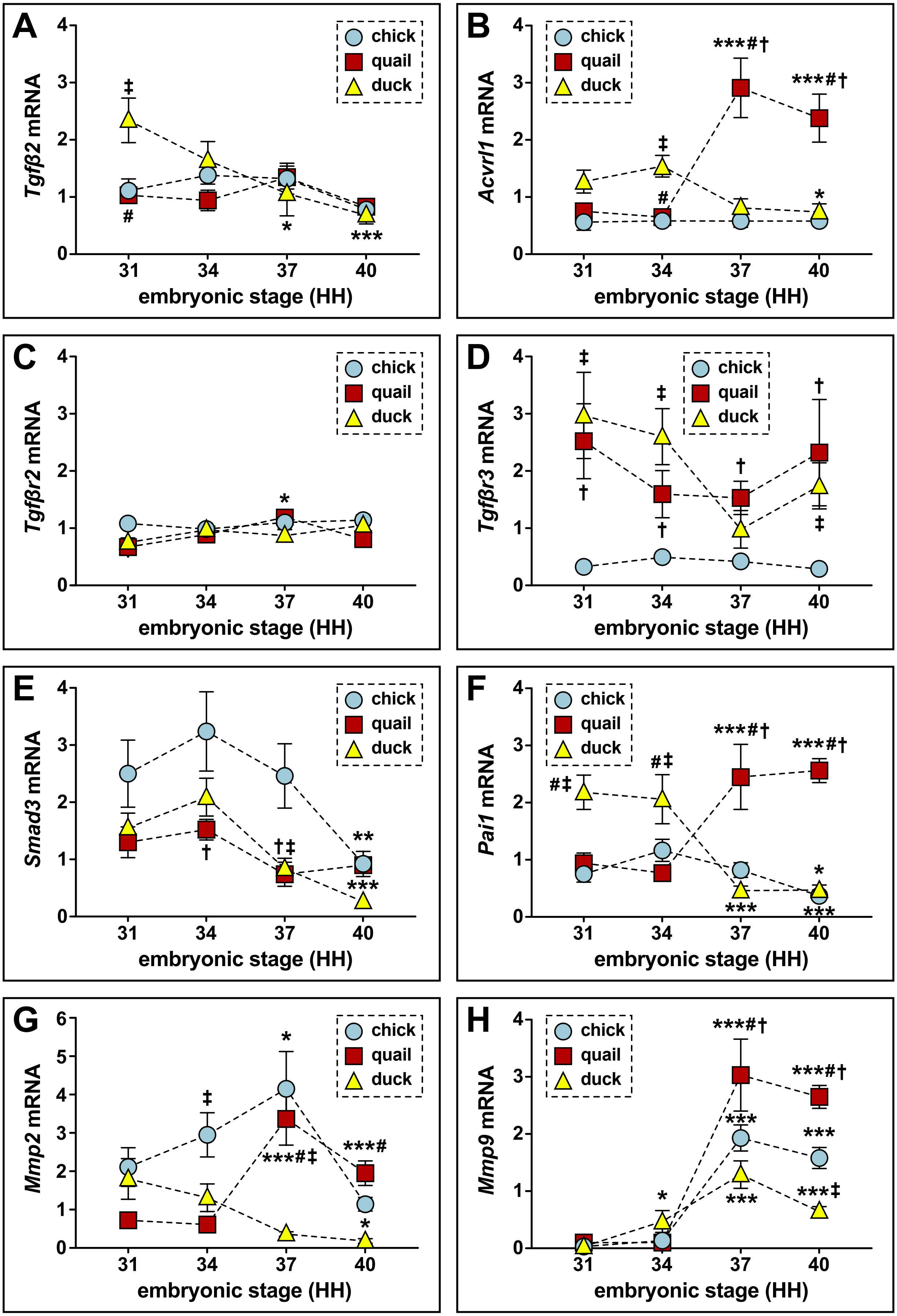


**SUPPLEMENTAL FIGURE 2**

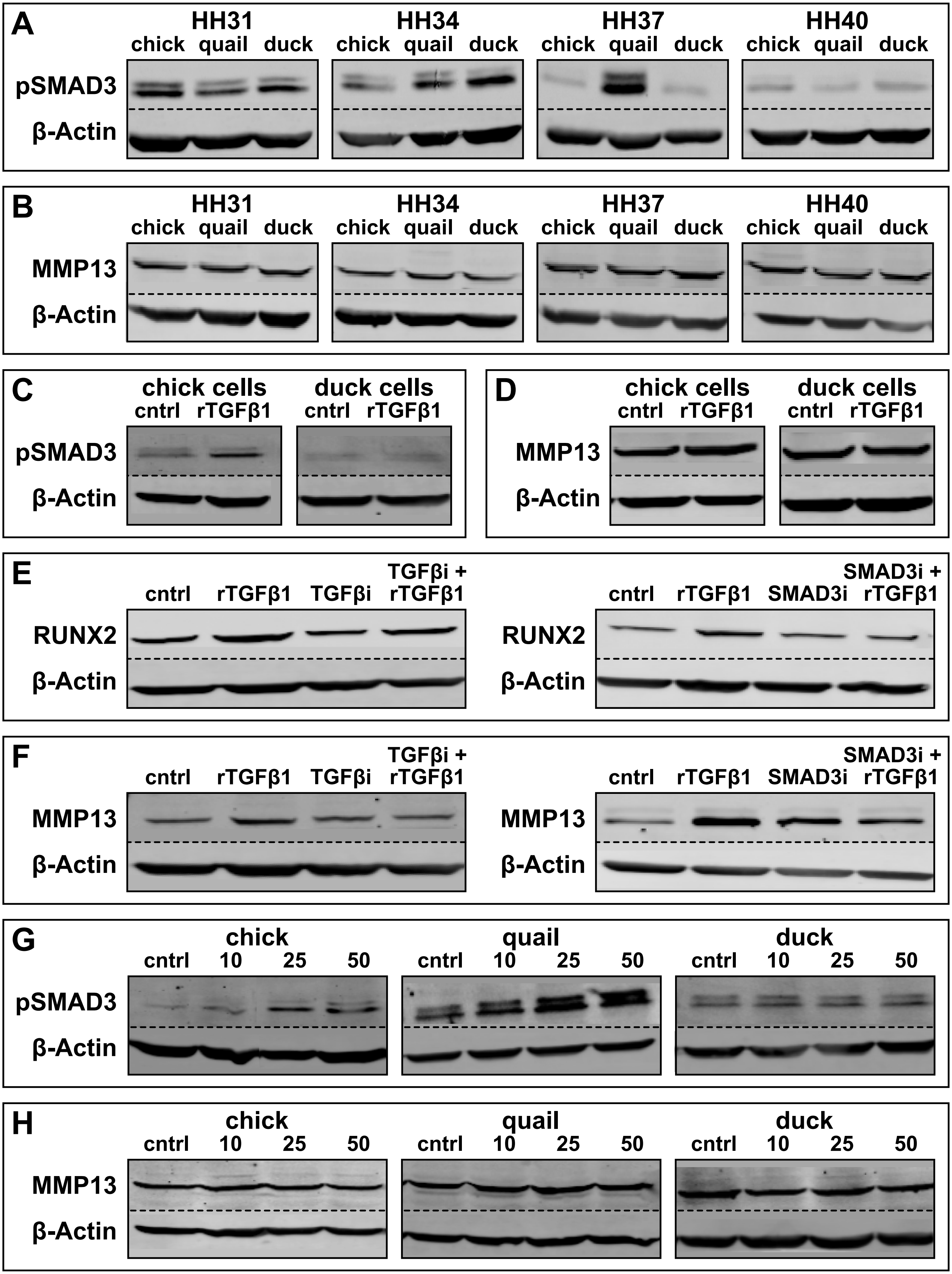


SUPPLEMENTAL FIGURE 3

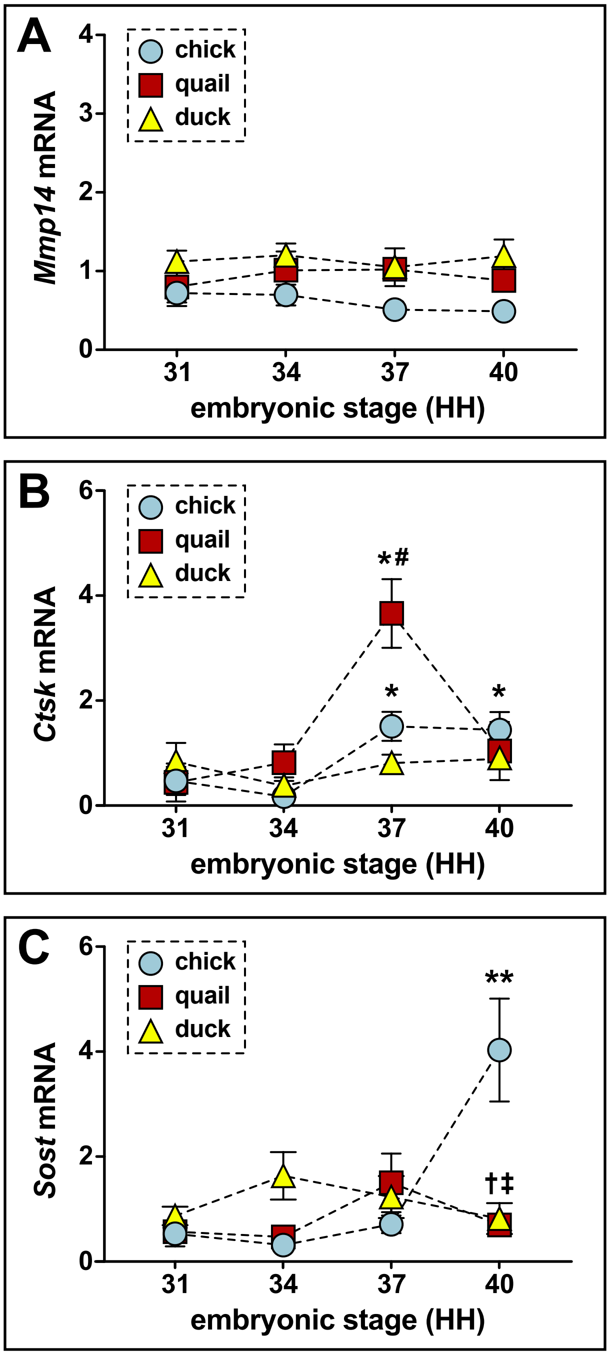


SUPPLEMENTAL FIGURE 4

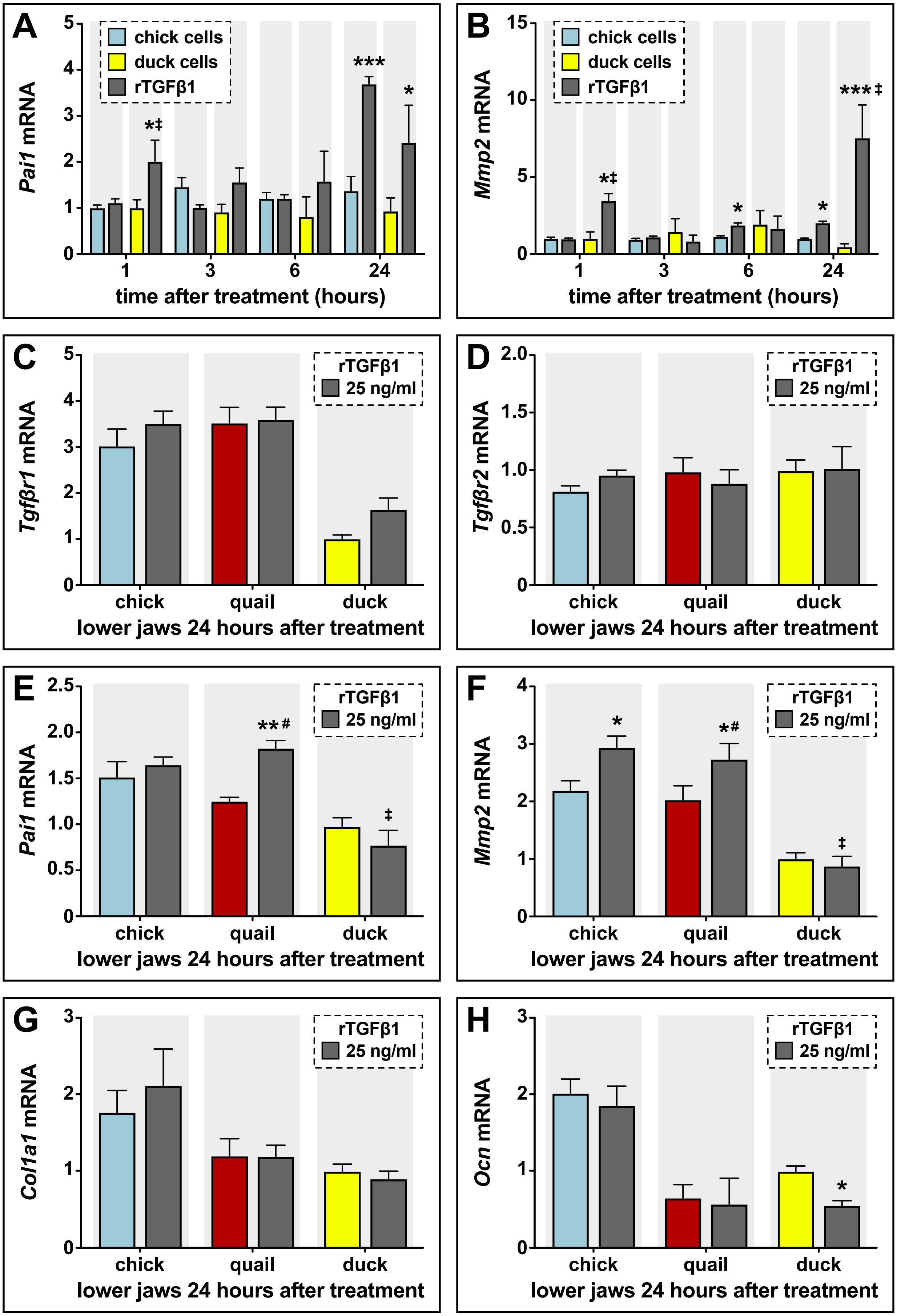


**SUPPLEMENTAL FIGURE 5**
