## Supplemental Tables for "Species-specific sensitivity to TGFβ signaling and changes to the Mmp13 promoter underlie avian jaw development and evolution"

**Supplemental Table 1.** *MMP13 antigen sequence*

| | Antigen Sequence |
|---|---|
| MMP13 antibody for chick, quail, duck. Affinity-purified peptide supplied polyclonal. Host strain: New Zealand rabbit. | MHHHHHHAPLHSKPQAVITFPGELLSAPSDVELAENY<br>LLRFGYIQEAEVRRSSKHVSLAKALRRMQKQLGLEET<br>GELDASTLEAMRAPRCGVPDVGGFLTFEDELKWDHM<br>DLTYRVMNYSPDLDRAVIDDAFRRAFKVWSDVTPLTF<br>TQIYSGEADIMIMFGSQEHGDGYPFDGKDGLLAHAFPP<br>GSGIQGDAHFDDDEFWTLGTGLEVKTRYGNANGASCH<br>FPFIFEGRSYSRCITEGRTDGMLWCATTASYDADKTYGF<br>CPSELLYTNGGNSDGSPCVFPFIFDGASYDTCTTDGRSD<br>GYRWCATTANFDQDKKYGFCPNRDTAAIGGNSQGDPC<br>VFPFTFLGQSYSARTSQGRQDGKLWCATTSNYDTDKK<br>WGFCPDPGYSIFLVAAH |

| **Supplemental Table 2.** *Primer sequences used for PCR and qPCR analysis* | |
|---|---|
| Gene | Sequence |
| Chick/quail/duck *18S* (sense) | 5'- GCGTGTGCCTACCCTACGCC -3' |
| Chick/quail/duck *18S* (antisense) | 5'- ACGCAAGCTTATGGCCCGCA -3' |
| Chick/quail/duck *Tgfβ1* (sense) | 5'- TCATGACATGAACCGGCC -3' |
| Chick/quail/duck *Tgfβ1* (antisense) | 5'- AGTCAATGTACAGCTGCC -3' |
| Chick/quail/duck *Tgfβ3* (sense) | 5'- AATGCACTGCTATCTCCTG -3' |
| Chick/quail/duck *Tgfβ3* (antisense) | 5'- CTTCAGCTTGCTCAGGATC -3' |
| Chick/quail/duck *Tgfβr1* (sense) | 5'- GTACATGGCACCTGAAGTTC -3' |
| Chick/quail/duck *Tgfβr1* (antisense) | 5'- GGCAACTGGTAATCTTCATG -3' |
| Chick/quail/duck *Smad2* (sense) | 5'- TTGCTGCTCTTCTGGCTC -3' |
| Chick/quail/duck *Smad2* (antisense) | 5'- TGAAGTTCAATCCAGCAAGG -3' |
| Chick/quail *Runx2 full length* (sense) | 5'- GCTGTGATGAAGAACCAGGT -3' |
| Duck *Runx2 full length* (sense) | 5'- GCCGTGATGAGGAACCAGGT -3' |
| Chick/quail/duck *Runx2 full length* (antisense) | 5'- TGGTATGTAGCGACTTGGGG -3' |
| Chick/quail/duck *Mmp13* (sense) | 5'- TTGGCTTGGAGGTGACAG -3' |
| Chick/quail/duck *Mmp13* (antisense) | 5'- GAACATCAGTGCTCCAGG -3' |
| Chick/quail/duck *Tgfβ2* (sense) | 5'- AATGCACTGCTATCTCCTG -3' |
| Chick/quail/duck *Tgfβ2* (antisense) | 5'- CTTCAGCTTGCTCAGGATC -3' |
| Chick/quail/duck *Acvrl1* (sense) | 5'- GTGAGACCGAGATCTACAAC -3' |
| Chick/quail/duck *Acvrl1* (antisense) | 5'- TCTGCAGGTAGTCGTAGAG -3' |
| Chick/quail/duck *Tgfβr2* (sense) | 5'- CTCACAAGAAGAGGAAGCTC -3' |
| Chick/quail/duck *Tgfβr2* (antisense) | 5'- CTGTGTTGTGGTTGATGTTG -3' |
| Chick/quail/duck *Tgfβr3* (sense) | 5'- ATGTGGATATGATCTTGGCC -3' |
| Chick/quail/duck *Tgfβr3* (antisense) | 5'- AGAGTGTCCAGGCCATAG -3' |
| Chick/quail/duck *Smad3* (sense) | 5'- CCTATGTCTCCAGCACAC -3' |
| Chick/quail/duck *Smad3* (antisense) | 5'- AAGCCATCCACGGTCATG -3' |
| Chick/quail/duck *Pai1* (sense) | 5'- TAGTGGCAGCGATAGATCC -3' |
| Chick/quail/duck *Pai1* (antisense) | 5'- ATCAGCAGGCGGATGTTC -3' |
| Chick/quail/duck *Mmp2* (sense) | 5'- GATGATGACCGCAAGTGG -3' |
| Chick/quail/duck *Mmp2* (antisense) | 5'- GGAAGTTCTTGGTGTAGGTG -3' |
| Chick/quail/duck *Mmp9* (sense) | 5'- GACACCGACAAGAAGTGG -3' |
| Chick/quail/duck *Mmp9* (antisense) | 5'- GAAGTCCTGGACGTAGCTG -3' |
| Chick *Runx2 promoter 1* (sense) | 5'- CGACAGGAACTATGGCATCA -3' |
| Quail/duck *Runx2 promoter 1* (sense) | 5'- CGAGAGGGACTATGGCATCA -3' |
| Chick/quail *Runx2 promoter 1 (*antisense) | 5'- CTGTTGGGCCACTACGACC -3' |
| Duck *Runx2 promoter 1* (antisense) | 5'- CTGTTGGGCTACTACGACC -3' |
| Chick/quail/duck *Runx2 promoter 2* (sense) | 5′- TTGTGGCTGTTGTGATGCG -3′ |
| Chick/quail *Runx2 promoter 2 (*antisense) | 5′- CTGTTGGGCCACTACGACC -3′ |
| Duck *Runx2 promoter 2* (antisense) | 5'- CTGTTGGGCTACTACGACC -3' |
| Chick/quail/duck *Runx2 exon 5* (sense) | 5′- GTCGCTACATACCACAGAGC -3′ |
| Chick/quail/duck *Runx2 exon 5 (*antisense) | 5′- TCGGGACCCCTACTCTCATA -3′ |
| Chick/quail/duck *Runx2 exon 4,6* (sense) | 5′- GTCGCTACATACCACAGAGC -3′ |
| Chick *Runx2 exon 4,6* (antisense) | 5′- GGAAGTCCTGTGCCTCGG -3′ |
| Quail/duck *Runx2 exon 4,6 (*antisense) | 5′- GGAAGTCCTGTGCCTCGA -3′ |
| Chick/quail/duck *Runx2 c-terminus 1* (sense) | 5′- GACTTCCAGCCATCACCGA -3′ |
| Chick/quail/duck *Runx2 c-terminus 1* (antisense) | 5′- AGTGAGTGGTTGCGGACATA -3′ |
| Chick/quail/duck *Runx2 c-terminus 2* (sense) | 5′- GACTTCCAGCCATCACCGA -3′ |
| Chick/quail *Runx2 c-terminus 2* (antisense) | 5′- GAAGTTGGTGGAGAAGTGGC -3′ |

| | |
|---|---|
| Duck *Runx2 c-terminus 2* (antisense) | 5′- GAAGTTGGTGGTGAAGTGGC -3′ |
| Chick/quail/duck *Ocn (sense)*<br>Chick/quail/duck *Ocn (antisense)* | 5′- TCACATTCAGCCTCTGCCG -3′<br>5′- GCTCACACACCTCTCGTTGG -3′ |
| Chick/quail *Col1a1* (sense)<br>Duck *Col1a1* (sense)<br>Chick/quail *Col1a1* (antisense)<br>Duck *Col1a1* (antisense) | 5′- AGAACAGCGTCGCCTACATG -3′<br>5′- TGCCTTCATCTGGTGCTCAC -3′<br>5′- CTCCAGTGTGACTCGTGCAG -3′<br>5′- GTGTCCTGGTCCATGTAGGC -3′ |
| Chick/quail/duck *Alp (sense)*<br>Chick/quail/duck *Alp* (antisense) | 5′- TCCACCAGCAGGAAGAAG -3′<br>5′- GGATATGGTGTATGAGCTGG -3′ |
| Chick *Mmp13 full length (sense)*<br>Chick *Mmp13 full length* (antisense) | 5′- ATGCAACCCAGACTTTCAGC -3′<br>5′- GGTAGTCAGTGCTTGTTCGC -3′ |
| Duck *Mmp13 full length (sense)*<br>Duck *Mmp13 full length* (antisense) | 5′- TCAGGGCTTCAGACTTCACA -3′<br>5′- AGCCTGCAACATGTCATTCA -3′ |
| Quail *Mmp13 full length (sense)*<br>Quail *Mmp13 full length* (antisense) | 5′- ACACATCAGGGCTTCAGACT -3′<br>5′- TGCAGCTTGTAACATGGCAC -3′ |
| Chick pTet *Mmp13 (sense)*<br><br>Quail pTet *Mmp13 (sense)*<br><br>Chick/quail pTet *Mmp13* (antisense)<br><br>Duck pTet *Mmp13 (sense)*<br><br>Duck pTet *Mmp13* (antisense) | 5′-ACCCTCGTAAAGCCGCCACCATGCAACCCAGACTTTCAGC -3′<br><br>5′-ACCCTCGTAAAGCCGCCACCATGCAACCAAGACTTTCAGC -3′<br><br>5′-GCCGCTTCACTTGTACTGCATCAGCACCAAAATAAGGAGT -3′<br><br>5′-ACCCTCGTAAAGCCGCCACCATGATGCAGTCAAGGCTTTC -3′<br><br>5′- GCCGCTTCACTTGTACTGCATCAGCACCAAAATATGGAGT -3′ |

| **Supplemental Table 3.** *Potential binding elements within the Mmp13 promoter predicter by JASPAR* | | | | | | |
|---|---|---|---|---|---|---|
| **Chick** | | | | | | |
| Transcription factor | Position | Absolute score | Relative score | Strand | JASPAR ID | Sequence |
| SMAD2::SMAD3 | 73-86 | 13.71 | 0.95 | + | MA1622.1 | GTGGTGACTCACTG |
| RUNX2 | 163-171 | 11.84 | 0.96 | + | MA0511.2 | AAACCACAG |
| RUNX2 | 189-197 | 9.40 | 0.93 | + | MA0511.2 | AGACCACAG |
| SMAD4 | 1046-1053 | 8.19 | 0.93 | - | MA1153.1 | AGTCTGGG |
| RUNX2 | 1108-1116 | 9.74 | 0.94 | - | MA0511.2 | AAACCTCAA |
| SMAD2::SMAD3 | 1252-1265 | 11.38 | 0.91 | + | MA1622.1 | TAAATGACTAATAC |
| SMAD3_1 | 1841-1857 | 11.21 | 0.90 | + | PB0060.1 | CTTCTCCAGACTGAACA |
| SMAD4 | 1846-1853 | 10.17 | 0.96 | - | MA1153.1 | AGTCTGGA |
| **Duck** | | | | | | |
| SMAD2::SMAD3 | 70-83 | 13.71 | 0.95 | + | MA1622.1 | GTGGTGACTCACTG |
| RUNX2 | 160-168 | 11.84 | 0.96 | + | MA0511.2 | AAACCACAG |
| RUNX2 | 485-493 | 7.37 | 0.91 | - | MA0511.2 | ACACCACAT |
| SMAD4 | 838-845 | 13.44 | 1.00 | - | MA1153.1 | TGTCTAGA |
| SMAD1 | 1173-1186 | 11.24 | 0.90 | + | UN0263.1 | ATTTTTATCTGTTT |
| RUNX2 | 1733-1741 | 11.84 | 0.96 | - | MA0511.2 | AAACCACAG |
| **Quail** | | | | | | |
| SMAD2::SMAD3 | 72-85 | 11.96 | 0.92 | - | MA1622.1 | ACGATGAGTCACCA |
| SMAD2::SMAD3 | 73-86 | 14.80 | 0.97 | + | MA1622.1 | CTGGTGACTCATCG |
| RUNX2 | 163-171 | 11.84 | 0.96 | + | MA0511.2 | AAACCACAG |
| RUNX2 | 869-877 | 10.94 | 0.95 | + | MA0511.2 | AAACCACAT |
| SMAD3_1 | 1038-1054 | 11.71 | 0.91 | + | PB0060.1 | TGAATTCAGACAAAACA |
| RUNX2 | 1089-1097 | 9.74 | 0.94 | - | MA0511.2 | AAACCTCAA |
| SMAD4 | 1783-1790 | 10.03 | 0.96 | - | MA1153.1 | GGTCTGGA |
